# *Gtf2i* and *Gtf2ird1* mutation are not sufficient to reproduce mouse phenotypes caused by the Williams Syndrome critical region

**DOI:** 10.1101/558544

**Authors:** Nathan Kopp, Katherine McCullough, Susan E. Maloney, Joseph D. Dougherty

## Abstract

Williams syndrome is a neurodevelopmental disorder caused by a 1.5-1.8Mbp deletion on chromosome 7q11.23, affecting the copy number of 26-28 genes. Phenotypes of Williams syndrome include cardiovascular problems, craniofacial dysmorphology, deficits in visual spatial cognition, and a characteristic hypersocial personality. There are still no genes in the region that have been consistently linked to the cognitive and behavioral phenotypes, although human studies and mouse models have led to the current hypothesis that the general transcription factor 2 I family of genes, *GTF2I* and *GTF2IRD1*, are responsible. Here we test the hypothesis that these two transcription factors are sufficient to reproduce the phenotypes that are caused by deletion of the Williams syndrome critical region (WSCR). We compare a new mouse model with loss of function mutations in both *Gtf2i* and *Gtf2ird1* to an established mouse model lacking the complete WSCR. We show that the complete deletion model has deficits across several behavioral domains including social communication, motor functioning, and conditioned fear that are not explained by loss of function mutations in *Gtf2i* and *Gtf2ird1*. Furthermore, transcriptome profiling of the hippocampus shows changes in synaptic genes in the complete deletion model that are not seen in the double mutants. Thus, we have thoroughly defined a set of molecular and behavioral consequences of complete WSCR deletion, and shown that other genes or combinations of genes are necessary to produce these phenotypic effects.

## Introduction

Contiguous gene disorders provide a unique opportunity to understand genetic contributions to human biology, as their well-defined genetic etiology delimits specific genomic regions strongly affecting particular phenotypes. Williams syndrome (WS; OMIM #194050) is caused by a 1.5-1.8Mbp deletion of 26-28 genes on chromosome 7q11.23 in the Williams syndrome critical region (WSCR). Williams syndrome is phenotypically characterized by supravalvular aortic stenosis, craniofacial dysmorphology, and a distinct cognitive profile consisting of intellectual disability, severe visual-spatial deficits, yet relatively strong language skills. Other common cognitive and behavioral difficulties include high levels of anxiety, specific phobias, and a characteristic hypersocial personality manifested as strong eye contact, indiscriminate social approach, and social disinhibition (see (1–3) for reviews). Despite increased social interest, individuals with Williams syndrome have difficulties with social awareness and social cognition (4, 5). In contrast, the reciprocal duplication results in dup7q11.23 syndrome (OMIM #609757), which presents with both similar and contrasting phenotypes to WS, such as high levels of anxiety yet less social interest (6). It is also associated with autism spectrum disorders (7). The recurrent deletion and duplications of chr7q11.23 indicate that one or more genes in this region are dose sensitive and have a large effect on human cognition as well as human social behavior.

Substantial efforts have been taken to understand which genes in the WSCR contribute to different aspects of the phenotype. Three approaches have driven advances in genotype-phenotype correlations in the WSCR: phenotyping individuals with atypical deletions in the region, human induced pluripotent stem cell models, and mouse models. Patients with atypical deletions have firmly connected haploinsufficiency of the elastin (*ELN*) gene with supravalvular aortic stenosis and other elastic tissue difficulties in WS (8, 9). However, human studies have not conclusively linked other genes to specific phenotypes. Three atypical deletions that span the *ELN* gene to the typical telomeric breakpoints showed the full spectrum of the WS phenotype, suggesting that most of the phenotypes are driven by the telomeric end of the deletion, which contains genes for two paralogous transcription factors *GTF2I* and *GTF2IRD1* (10, 11). Indeed, most of the atypical deletions that have been reported that delete the centromeric end of the region and don’t affect the copy number of *GTF2I* and GTF2IRD1, show mild to none of the characteristic facial features or cognitive and behavioral phenotypes of WS (12–20). While there are contrasting examples of deletions that spare *GTF2I* and still have mild facial characteristics of WS, lower IQ, and the overfriendly social phenotype (12, 21), the preponderance of evidence from these rare partial deletions have led to the dominant hypothesis being that *GTF2I* and *GTF2IRD1* mutation are necessary to cause the full extent of the social, craniofacial, visual-spatial and anxiety phenotypes. However, there are limitations to these human studies, primarily due to the rarity of partial deletions. First, because of the variable expressivity of the phenotypes even in typical WS, it can be difficult to confidently interpret any phenotypic deviation in the rare partial deletions (4, 5, 22). Second, given the rarity of WS and partial deletions, and lack of relevant primary tissue samples, it is challenging to link genetic alterations to the specific downstream molecular and cellular changes that could mediate the organismal phenotypes.

To overcome this second barrier, researchers have turned to using patient induced pluripotent stem cells to study the effects of the WSCR deletion and duplication on different disease relevant cell types (23–27). While linking molecular changes to organismal behavior is not possible with cell lines, this approach is amenable to studying cellular and molecular phenotypes, such as changes to the transcriptome and cellular physiology. By studying differentiated neural precursor cells from an individual with a typical WS deletion and an individual with an atypical deletion that spares the copy number of the *FZD9* gene, Chailangkarn et al. (23) showed that *FZD9* is responsible for some of the cellular phenotypes, such as increased apoptosis and morphological changes. Lalli et al. (25) used a similar approach to show that knocking down the *BAZ1B* gene in differentiated neurons was sufficient to reproduce the transcriptional differences and deficits in differentiation that were observed in WS differentiated neurons. Finally, Adamo et al. (24) studied the effects of *GTF2I* on iPSCs from typical WS deletions, dup7q11.23, and typical controls. By overexpressing and knocking down *GTF2I* in the three genotypes, they showed that *GTF2I* was responsible for 10-20% of the transcriptional changes. Overall, using iPSCs from patients with WS has highlighted a role for both the *GTF2I* family and other less appreciated genes in the molecular consequences of the WSCR mutation. This suggested the possibility that several genes may play a role in the cognitive phenotypes and *GTF2I* alone may not be sufficient for all neural molecular changes and hence cognitive phenotypes. However, iPSC studies face the limitation that they cannot be used to model whole organismal effects like anxiety or social behavior. Further, while some cellular and molecular phenotypes can be evaluated, both gene expression and cellular physiology using *in vitro* differentiation systems do not perfectly reflect the phenotype of mature neural cells, fully integrated into a functioning or dysfunctioning brain.

Mouse models have been used to link genes in WSCR to specific molecular and cellular phenotypes, as well as to the functioning of conserved organismal behavioral circuits that could be related to human cognitive phenotypes. Mouse models are particularly suitable because a region on mouse chromosome five is syntenic to the WSCR, enabling models of corresponding large deletions, including a mouse line with a complete deletion (CD) of the WSCR genes that shows both behavioral disruptions and altered neuronal morphology (28). In addition, a key advantage over human partial deletions is that researchers can easily manipulate the mouse genome to delete targeted subsets of genes in the locus, and generate large numbers of animals with identical partial mutations, enabling statistical analyses to overcome variable expressivity. For example, there are mouse models of large deletions that show that genes in the distal and proximal half of the region may contribute to separate and overlapping phenotypes (29). Likewise, many single gene knockouts exist that show some phenotypic similarities to the human syndrome, though a limitation is that some of these studies model full homozygous loss of function, rather than a hemizygous decrease in gene dose. Nonetheless, specifically for *Gtf2ird1* (30–32) and *Gtf2i* (33–35), multiple mouse models of either gene have shown extensive behavioral deficits including social and anxiety-like behaviors, some of which present contrasting evidence. However, each of these studies has been conducted in isolation, by different labs, with fairly different phenotyping assays, making it difficult to directly compare findings to other mouse models of WS.

Mouse models uniquely enable a direct way to test the sufficiency of individual mutations to recreate the organismal phenotypes detected when the entirety of the WSCR is deleted. By crossing different mutant lines together, we can create genotypes unavailable in human studies and conduct a well-powered and controlled study to directly test if specific gene mutations are sufficient to reproduce particular phenotypes of the full deletion. Since both human and mouse literature suggest that *GTF2IRD1* and *GTF2I* each contribute to the molecular, cognitive, and social phenotypes, we set out here to test if loss of function of both of these genes is sufficient to recapitulate the phenotypes of the entire WSCR deletion at both the molecular and behavioral circuit levels, or if instead, as hinted by the iPSC studies and other human mutations, other or more genes may be involved. Using CRISPR/Cas9 we generated a new mouse line that has loss of function mutations in both *Gtf2i* and *Gtf2ird1* on the same chromosome. We then crossed them to the CD full deletion model to directly compare behavior and transcriptomes of the *Gtf2i*/*Gtf2ird1* mutants to both WT and CD littermates. Examining both previously defined and newly characterized behavioral and molecular disruptions, we demonstrate that mutation of these two genes is not sufficient to replicate *any* of the CD phenotypes. In contrast to a dominant hypothesis arising from human partial deletions, this study provides strong evidence that *Gtf2i*/*Gtf2ird1* mutation alone may not be responsible for key WS cognitive and behavioral phenotypes.

## Results

### Generation and validation of Gtf2i and Gtf2ird1 loss of function mutation on the same chromosome

Prior work from comparing phenotypes of humans with partial deletions of the WSCR highlighted *GTF2I* and *GTF2IRD1* as likely involved in cognitive phenotypes in WS (10, 13, 20). Likewise, single gene mutant mouse models of both genes showed that each may contribute to relevant phenotypes (30–33, 36). We wanted to test if heterozygous loss of function mutants of both *Gtf2i* and *Gtf2ird1* are sufficient to replicate the phenotypes that are caused when animals are hemizygous for the entire WSCR (Figure 1A).

**Figure 1.**
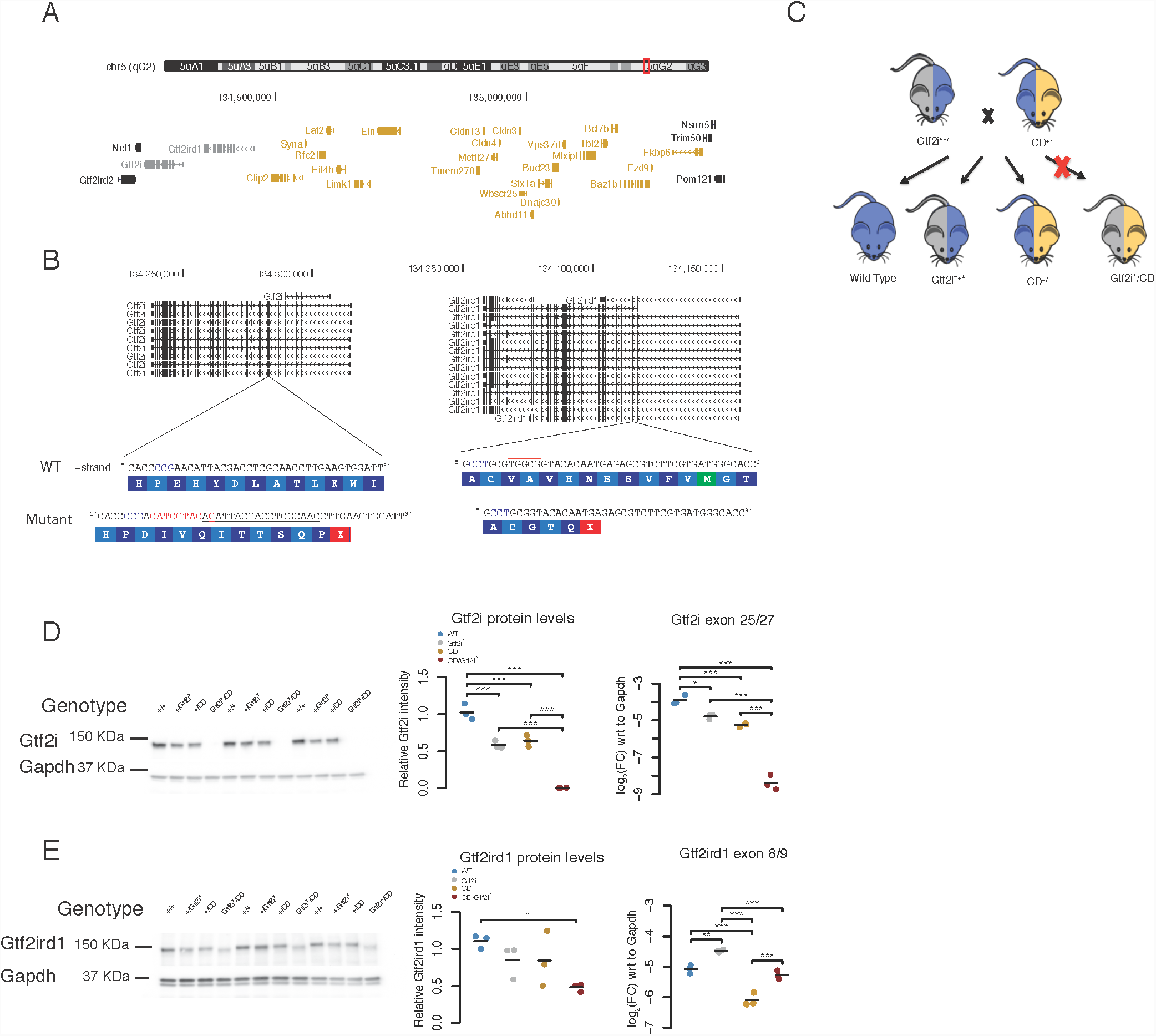
Generation of double mutant GTF2I*\** model. **A** Schematic of the syntenic WSCR in mouse on chromosome 5. The two transcription factors being tested here are highlighted in grey and the genes that are deleted in the CD animals are highlighted in yellow. **B** Gene models of *Gtf2i* and *Gtf2ird1* showing the multiple isoforms of each gene. The WT sequences with the gRNA target underlined and the PAM highlighted in blue with the mutant sequences below along with the corresponding amino acid sequence. **C** Breeding scheme for the behavior tasks **D.** E13.5 whole brain Gtf2i western and qPCR of *Gtf2i\** x CD. Gtf2i protein and transcript are similarly reduced in the *Gtf2i\** and CD animals. **E** E13.5 whole brain Gtf2ird1 western and qPCR of *Gtf2i\** x CD. Gtf2ird1 protein is slightly reduced in the *Gtf2i*/*CD brain compared to WT. *Gtf2ird1* transcript is increased in the *Gtf2i\** genotype, decreased in the CD genotype, and returns to WT levels in *Gtf2i*/*CD genotype. * p < 0.05, ** p < 0.01, *** p < 0.001

Therefore, to test the sufficiency of these genes, we generated a mutant of *Gtf2i* and *Gtf2ird1* genes on the same chromosome using CRIPSR/Cas9. Two gRNAs were designed to target constitutive exons of *Gtf2i* or *Gtf2ird1* (Figure 1B) and were co-injected with Cas9 mRNA into the eggs of the FVB strain. Of the 57 pups born we detected 21 editing events using the T7 endonuclease assay. From these animals PCR amplicons around each targeted site were deeply sequenced and mutations were characterized via manual inspection of the reads in IGV. Of the founders there were five that only had mutations in *Gtf2i*, five with mutations only in *Gtf2ird1*, and 15 that had mutations in both genes (Supplemental Figure 1A). Most founders had more than one allele within a gene indicating high rates of mosaicism (60%, 15/25 mice). Breeding a selection of the mosaic founders to WT animals revealed that some of the founders were mosaic in the germline as well (40%, 4/10 mice), with one founder transmitting three different alleles.

To test if haploinsufficiency of both *Gtf2i* and *Gtf2ird1* is sufficient to replicate the phenotype of hemizygosity of the entire WSCR, we moved forward with characterizing a mouse line that has a G > C polymorphism followed by an eight base pair insertion in exon five of *Gtf2i* and a five base pair deletion in exon three of *Gtf2ird1*; these will be referred to as the *Gtf2i\** mouse line (Figure 1B). These mutations are inherited together indicating that they are on the same chromosome. The mutations cause frameshifts and introduce premature stop codons in early constitutive exons (Figure 1B), and were thus expected to trigger nonsense mediated decay and lead to loss-of-function alleles, mimicking the effective gene dosage of WSCR region deletions for these two genes.

We first performed RT-qPCR and western blots to confirm the effects of the frameshift mutations at the transcript and protein levels in embryonic day 13.5 (E13.5) littermates that were WT, heterozygous, and homozygous mutant at the locus. We used E13.5 brains for two reasons 1) homozygosity of *Gtf2i* null mutants is embryonic lethal (33, 37) and 2) both Gtf2i and Gtf2ird1 proteins are more highly expressed during embryonic time points in the brain, with undetectable levels of Gtf2ird1 in the WT adult mouse brain (Supplemental Figure 1B and C).

The frameshift mutation in exon five of *Gtf2i* reduced the amount of transcript detected by qPCR, consistent with nonsense mediated decay. This mutation led to a 50% decrease of the protein in heterozygous animals and no protein in homozygous mutants (Supplemental Figure 1D). Indeed we were not able to recover pups that were homozygous for the *Gtf2i\** mutations after birth, but we were able to harvest homozygous embryos up to E15.5. The embryos had exencephaly consistent with other *Gtf2i* mouse models (33, 37).

In contrast, the frameshift mutations in exon three of *Gtf2ird1* increased the amount of transcript, as expected. Increases in transcript of *Gtf2ird1* due to a loss of function mutation have been described in another *Gtf2ird1* mouse model, and both EMSA and luciferase reporter assays indicated that Gtf2ird1 protein represses the transcription of the *Gtf2ird1* gene (38). The increase in transcript was commensurate with the dosage of the mutation (Supplemental Figure 1E). However, we saw that the protein levels in our mutants did not change with dosage of the mutation and did not follow the trend of the transcript (Supplemental Figure 1E).

Production of detectable protein after the frameshift was surprising, especially since the increased *Gtf2ird1* mRNA levels were indeed consistent with prior studies of loss of functional Gtf2ird1 protein, so we investigated this phenomena further. We noticed that the homozygous Gtf2ird1 protein bands looked slightly shifted in the western blots. This lead us to hypothesize that there could be a translation reinitiation event at the methionine in exon three downstream of the frameshift mutation in a different open reading frame (Supplemental Figure 1F). In another targeted mutation of *Gtf2ird1*, where the entire exon two, which contains the conical start codon, was removed, the authors noted that there was still three percent of protein being made, and the product that was made was similarly shifted (38). From our mutation we would expect a 65aa N-terminal truncation, which corresponds to a 7KDa difference between WT. We ran a lower percentage PAGE gel to get better separation between WT and homozygous animals and we saw a slight shift, suggesting that there was reinitiation of translation at methionine-65 in a different open reading frame (Supplemental Figure 1G). This was indicative of the loss of the N-terminal end of the protein, which contains a leucine zipper that is thought to be important in DNA binding (38). This is consistent with the mRNA evidence that the allele is loss of function.

We therefore tested the hypothesis that we had abolished the DNA binding capacity of the truncated protein, to confirm loss of function. We performed ChIP-qPCR and pulled down DNA bound to Gtf2ird1 protein and then amplified the promoter region of *Gtf2ird1,* which has previously been shown to be bound by the Gtf2ird1 protein. We compared this to two off-target regions in the genome near *Bdnf* and *Pcbp3*. We performed this experiment in E13.5 brains of WT and homozygous *Gtf2i\** embryos. There was a 15-20 fold enrichment of the on target *Gtf2ird1* promoter region compared to the off target regions in the WT animals, while the truncated protein did not show any enrichment (Supplemental Figure 1H and 1I). This suggested that while a truncated protein was still being made it did not have the DNA binding functionality of the WT protein. This indicated that the frameshift mutation in exon three of *Gtf2ird1* was a loss-of-function mutation and provided evidence that the N-terminal end of the protein, which contains a leucine zipper, is necessary for DNA binding. Thus, we confirmed we had generated a mouse line with loss of function alleles on the same chromosome for these *Gtf2i*\* genes.

To test the sufficiency of mutation in these two transcription factors to replicate phenotypes observed by deleting the entire WSCR, we crossed the *Gtf2i\** mutant to the CD mouse (Figure 1C), which is hemizygous from exon five of *Gtf2i* to *Fkbp6* (Figure 1A). The *Gtf2i\** mutants were generated on the FVB/AntJ background, whereas the CD mice were generated on the C57BL/6J background. Therefore, we only used the first generation from this cross for all experiments to ensure all mice had the same genetic background. As above, we assessed the transcript and protein levels of genotypes from this cross to confirm loss of function. Again, the CD/*Gtf2i\** genotype was embryonic lethal, but we did observe that genotype up to E15.5. The levels of *Gtf2i* transcript and protein were similar between CD heterozygous and *Gtf2i\** heterozygous animals (Figure 1D). The levels of *Gtf2ird1* transcript increased in *Gtf2i\** animals similar to what was seen in *Gtf2i\** heterozygous animals on the pure FVB/AntJ background. In contrast, the CD heterozygous animals had decreased levels of *Gtf2ird1* transcript. In the CD/*Gtf2i\** animals the level of transcript returned to WT levels. Again, the levels of *Gtf2ird1* transcript were not reflected in the protein levels. We saw a trend to similar slight decreases in protein levels in the both heterozygous genotypes; however, they were not significantly different from WT levels. This was interesting because in the CD animals were missing one entire copy of this gene, opposed to a frameshift mutation. This also suggested that the frameshift mutation in exon three of *Gtf2ird1* did affect the amount of protein being made, but not drastically. We did see a significant decrease in protein levels (60% of WT) in the CD/*Gtf2i\** genotype (Figure 1E). Again suggesting that the frameshift mutation was decreasing the levels of protein.

### Gtf2i* mutation is not sufficient to reproduce WSCR-mediated alterations of vocal communication

We next tested if haploinsufficiency for both genes would recapitulate behavioral phenotypes seen in mice hemizygous for the entire WSCR (CD mice) (Table 1). Since single gene knockout studies of both *Gtf2i* and *Gtf2ird1*, and larger deletion models showed evidence for disrupted social behavior we wanted to directly compare the effects of *Gtf2i\** haploinsufficiency to the effects of hemizygosity of the entire WSCR on social behavior.

**Table 1:**
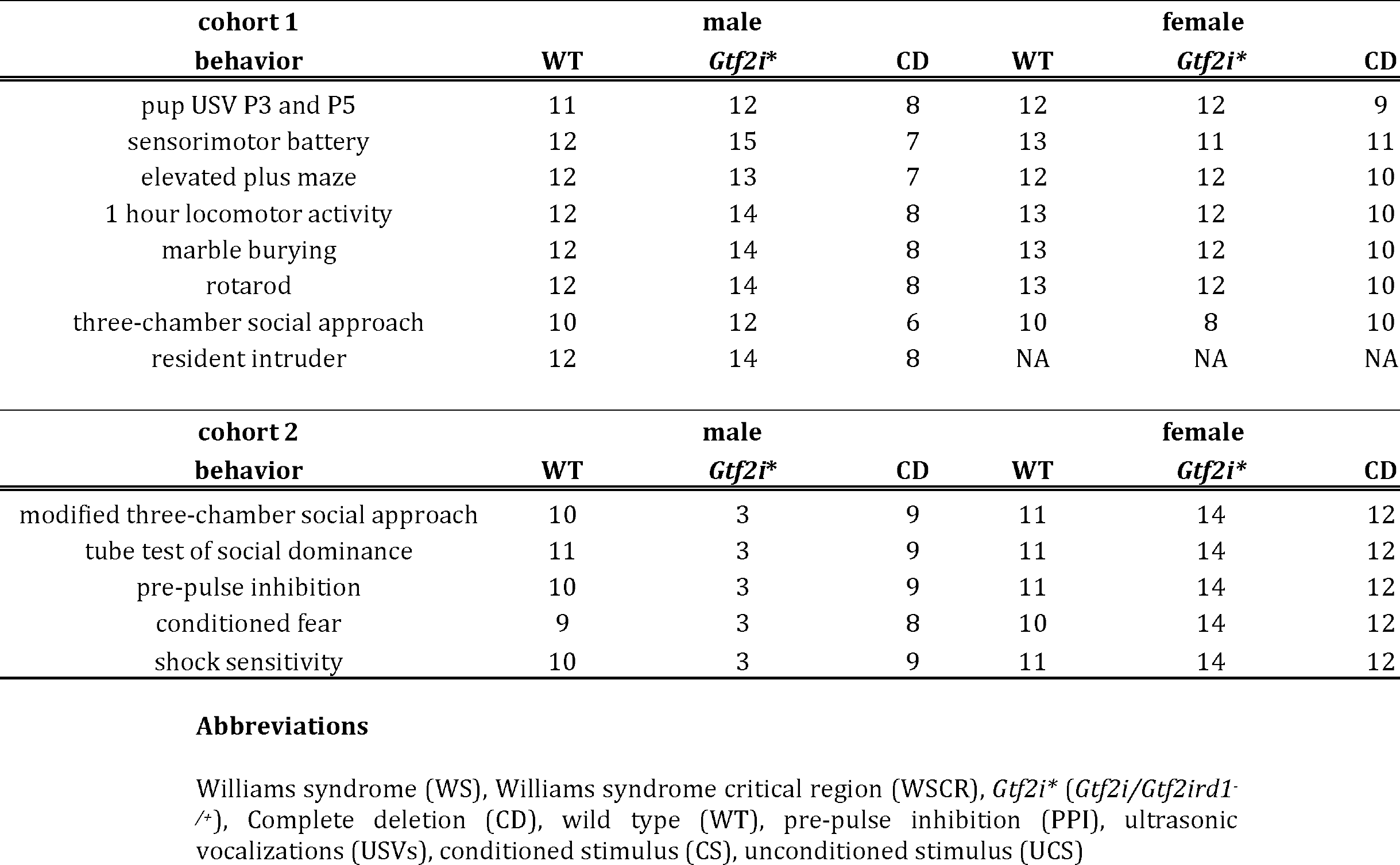
Behavior cohorts.

We first measured maternal separation induced ultrasonic vocalizations (USVs) in postnatal day three and postnatal day five pups. This is a form of developmental communication and was shown to be increased in mice that had three or four copies of *Gtf2i* compared to mice with normal copy number or only one functional copy (34). We saw a significant effect of day (F_1,116.00_=5.43, p=0.021) and genotype on the call rate (F_2,60.7_= 6.09, p=0.004), as well as a genotype by day interaction (F_2,61.64_=6.80, p=0.002). Post hoc analysis within day showed that on day five CD mice made fewer calls than WT littermates (p<0.001) and *Gtf2i\** mutant littermates (p=0.045) (Figure 2A). We included the weight of the mouse as a covariate to make sure the decrease in call number was not due to differences in weight. We saw that weight has a trending effect (F_1,75.48_=3.95, p =0.05), but the day by genotype interaction term remained significant.

**Figure 2.**
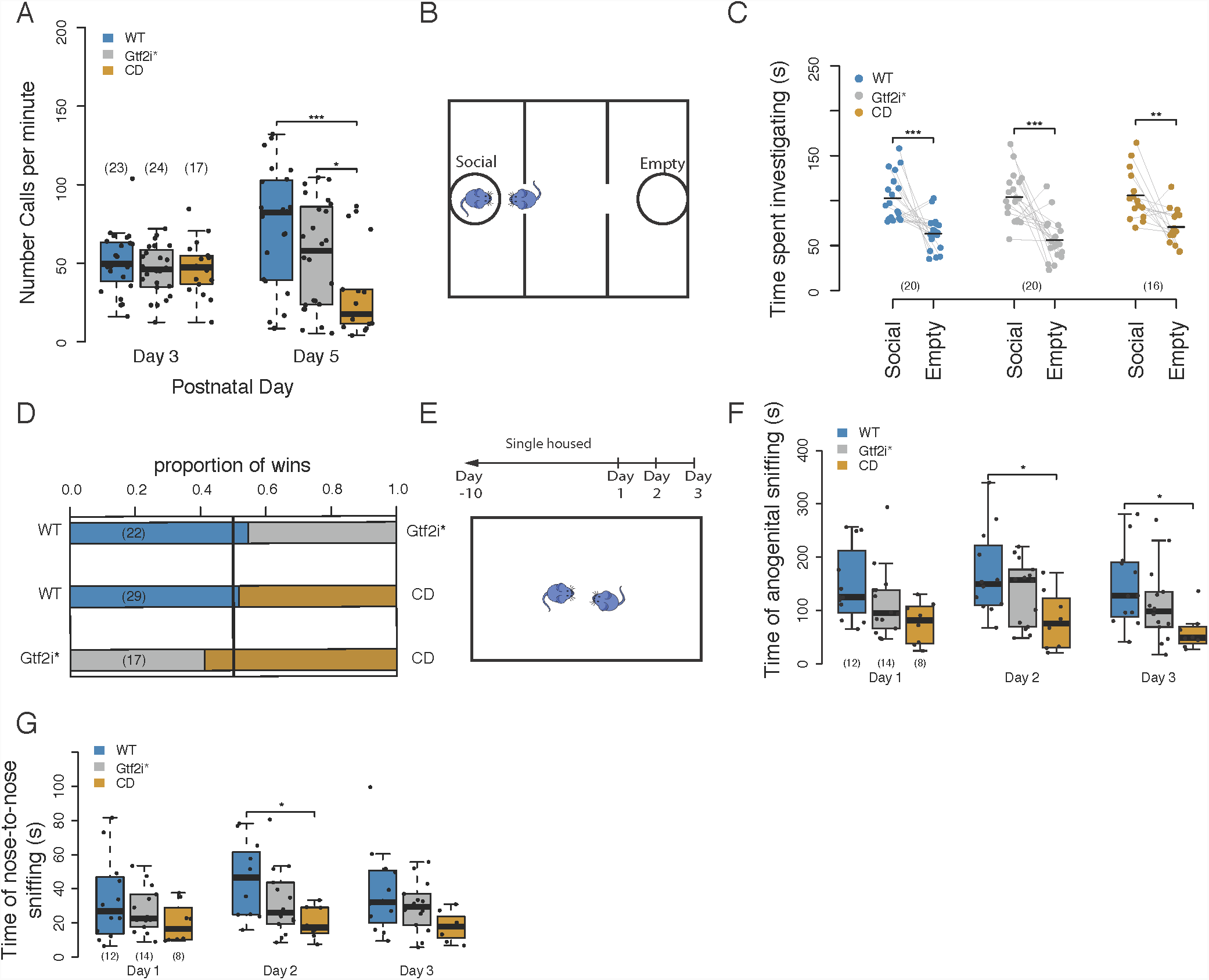
CD mice have deficits in ultrasonic vocalizations and decreased social investigation. **A** Callrate across two days shows that on postnatal day 5 CD animals produce fewer ultrasonic vocalizations than either WT or *Gtf2i\** littermates. **B** Schematic of the three-chamber social approach task. **C** All genotypes show preference for social stimulus in three-chamber social approach assay. **D** *Gtf2i\** and CD animals show similar dominance behavior to WT animals in the tube test for social dominance. **E** Schematic of the resident intruder paradigm. **F** CD animals show decreased time engaged in anogential sniffing in resident intruder task. **G** CD animals show decreased time engaged in nose-to-nose sniffing in resident intruder task. * p < 0.05, ** p < 0.01, *** p < 0.001 Sample sizes are shown as numbers in parentheses

We also observed differences in the temporal and spectral features of the calls. There was a significant effect of genotype on pause length between bouts (F_2,60_=11.9069, p=4.31e-5), with CD mice exhibiting longer pauses on day five compared to WT mice (p=0.0004) and *Gtf2i\** mice (p=0.0014); this is correlated with fewer calls produced by CD animals (Supplemental Figure 2A). There was a also significant genotype by day interaction for the duration of a call bout (F_2,61_=7.26, p=0.001), with CD mice exhibiting a shorter duration on day five compared to WT (p=0.046) (Supplemental Figure 2B). Overall, our study of vocalization provides evidence that *Gtf2i* and *Gtf2ird1* mutation alone are not sufficient to produce a CD-like deficit in this behavior.

Maternal-separation induced USVs are only produced during a transient period of development from postnatal day three to postnatal day 10, peaking at postnatal day seven and postnatal day nine in FVB/AntJ and C57BL/6J strains, respectively (39). Therefore the alteration in the CD animals could reflect an overall shift in developmental trajectory. To assess this, we checked weight gain and developmental milestones in our cohorts. No differences in developmental weights were observed between genotypes. The detachment of the pinnae at postnatal day five, a physical milestone, was similar across all genotypes (χ^2^=2.593, p=0.4628, Supplemental Table 1).. However, there were weight deficits in CD animals in adulthood (Supplemental Figure 2C). There was a significant effect of day on weight (F_4,240_=1610.9, p < 2.2e-16), a significant effect of genotype (F_2,60_=7.2059, p=0.001568), and a significant day by genotype interaction (F_8,240_=6.9258, p=3.332e-8). These data suggest that gross developmental delay in CD animals does not explain the observed communication deficit.

### Gtf2i* mutation is not sufficient to reproduce WSCR-mediated alterations of social behaviour

We went on to test adult social behaviors. We first applied the standard three chamber social approach, which has not been reported in CD mice. In this task the mice are allowed to freely explore an apparatus with three chambers: a center chamber, a social chamber that contains a cup with a sex and age-matched mouse, and an empty chamber that only contains an empty cup (Figure 2B). This test measures the voluntary social approach of mice. We saw the expected preference for the social stimulus across all mice (F_1,53_=83.2013, p=1.894×10^−12^), with no impact of genotype (F_2,53_=1.1516, p=0.3239) or genotype by stimulus interaction (F_2,53_=0.5845, p=0.5609). Post hoc comparisons within genotypes confirmed that all genotypes spent significantly more time investigating the social stimulus than the empty cup (WT p <0.001; *Gtf2i\** p < 0.001; CD p=0.00456; Figure 2C). Thus, sociability as measured in this task is not sensitive enough to discern a hypersocial phenotype in these animals.

In a test for social novelty, a novel stranger mouse was then placed in the empty cup. All genotypes showed the expected preference for the novel stimulus animal (F_1,53_=50.3816, p=3.137×10^−9^), again with no effect of genotype (F_2,53_=1.3948, p=0.2568) or genotype by stimulus interaction (F_2,53_=0.5642, p=0.5722). Post hoc comparisons showed that all the genotypes spent significantly more time investigating the novel stimulus (WT p < 0.001; *Gtf2i\** p =0.00321; CD p=0.0012; Supplemental Figure 2D). Additionally in this task, we did notice a significant effect of genotype on overall distance traveled (F_2,53_=3.98, p0.024) with the *Gtf2i\** mutants traveling further distance than the WT animals in the sociability trial (p=0.0305; Supplemental Figure 2E), and a corresponding trend during the social novelty trial (F2,53=2.87, p=0.115). This suggests that the double mutants have a slight hyperactive phenotype in this task that is not seen in the CD mutants.

Previous reports on social phenotypes in mouse models of WS have described a lack of habituation to a social stimulus. To test this we repeated the three-chamber social approach task in a new cohort of animals with an extended sociability trial to test if the *Gtf2i\** mutants or the CD animals showed the preference for the social stimulus after the prolonged amount of time. Similar to the classic three-chamber results we saw a significant effect of the social stimulus in the first five minutes (F_1,56_=19.3683, p=4.891e-5), there was a trend of a genotype effect (F_2,56_=3.098, p=0.053) and no interaction (F_2,56_=0.4650, p=0.6350). Interestingly, we observed a significant preference for the social chamber in the WT and *Gtf2i\** mutants, but the CD animals only trended in this direction (Supplemental Figure 2F). To determine if the CD mutants do indeed maintain a prolonged social interest compared to WT littermates, we examined the last five minutes of the 30 minute sociability trial. While there was a significant effect of stimulus (F_1,56_=4.82, p=0.03), there was still no effect of genotype (F_2,56_=0.0523, p=0.949) or an interaction (F_2,56_=0.454, p=0.637). In fact, the significant effect of chamber was driven by the proportion of animals investigating the novel empty cup more than the social stimulus (Supplemental Figure 2G). These data lead us to conclude that the double mutants and CD animals show a WT-like habituation to social stimulus in this task.

We also tested social dominance in the tube test in these mice. Previous studies using partial deletions of the WSCR showed that the proximal deletion which contains *Gtf2i* and *Gtf2ird1* as well as deletions of both the proximal and distal regions in mice resulted in different win/loss ratios than WT mice and mice lacking just the distal end of the WSCR (29). In contrast, here, the *Gtf2i\** and CD animals did not exhibit dominance behavior different than chance would predict (WT vs *Gtf2i\** p=0.8318, WT vs CD p=1). *Gtf2i\** and CD animals also had similar proportions of wins when paired together (*Gtf2i\** vs CD p=0.6291) (Figure 2D).

The contrasts in our findings with those reported in prior papers could be due to differences in background strain. Different inbred mouse strains show different dominance behavior (40), and other phenotypes, such as craniofacial morphology in WS models has been shown to be strain dependent (13, 30, 41). We tested the effects of the background strain of the *Gtf2i\** and CD models by performing the same task on the respective background of each line and comparing them to their WT littermates. Thi showed that the *Gtf2i\** mutants had a WT-like phenotype while the CD mice had a submissive phenotype with significantly more losses to WT littermates (Supplemental Figure 2H). Thus, the submissive phenotype of the CD allele is dependent on strain which is not observed in the *Gtf2i*\* mutants.

Finally, we tested the male mice in a resident-intruder paradigm. In this task, male mice were singly housed for 10 days to establish their territory and, in a series of three test days, novel WT C57BL/6J animals were introduced into their territories as intruders. This task measures both social interactions and bouts of aggression between two freely moving animals (Figure 2E). In our study, only one mouse showed aggressive behavior towards the intruder mouse, so we did not further quantify this behavior. Assessment of the social interactions showed a significant main effect of genotype (F_2,31_=5.241, p=0.011) with no effect of day (F_2,62_=2.470, p=0.093) or day by genotyping interaction (F_4,62_=0.1095, p=0.978). Post hoc tests within each day showed that the CD animals spent less total time on day two (p=0.0248) and day three (p=0.0318) engaged in anogenital sniffing compared to the WT animals (Figure 2F). These differences could not be explained by differences in total activity levels between the genotypes (F_2,31_=1.399, p=0.262; Supplemental Figure 2I). The decrease in total time spent in anogenital sniffing was driven by a shorter average bout time (F_2,31_=5.852, p=0.007, Supplemental Figure 2J) and not the number of times the animals initiated the sniffing behavior (F_2,31_=2.7961, p=0.0765; Supplemental Figure 2K). The same differences also held for nose-to-nose sniffing (Figure 2G). There was a significant effect of genotype (F_2,31_= 3.737, p=0.0352) and no effect of day (F_2,62_=3.01, p=0.056) or day by genotype interaction (F_4,62_=0.8156, p=0.520). Post hoc analysis showed that on day two the CD animals participated in nose-to-nose sniffing significantly less than the WT animals (p=0.0160), while the trend was present in the other days but was not significant. These results indicated that some aspect of social behavior was disrupted in these animals and *Gtf2i\** mutants could not recapitulate the full CD phenotype. While we predicted that the WS models would show increased social interest similar to the human condition, individuals with WS have difficulties with other aspects of social behavior, such as social cognition and social awareness (4, 5), which may be reflected in these data.

### Gtf2i* mutation is not sufficient to reproduce WSCR mediated alterations of motor behaviour

Along with a characteristic social behavior, WS also presents with other cognitive phenotypes including poor coordination, increased anxiety, specific phobias, repetitive behaviors, and mild intellectual impairment (42). Human studies and mouse models have suggested that *GTF2I* and *GTF2IRD1* contribute in aspects of the visual-spatial deficits and other cognitive phenotypes (17, 20). These genes are also highly expressed in the cerebellum, which could contribute to the coordination problems (43, 44). Therefore, we next tested if CD mice had any motor phenotypes and if haploinsufficiency of these two transcription factors were sufficient to reproduce any deficits.

We performed a sensorimotor battery to assess balance, motor coordination and strength in mutants and WT littermates. All genotypes were similar in the time to initiate walking, and reach the top of a 60 degree inclined screen or a 90 degree inclined screen. All genotypes were able to hang onto an inverted screen for the same amount of time (Supplemental Figure 3A-D). CD animals were significantly quicker on turning around on a pole and quicker to get off of the pole than WT animals (Supplemental Figure 3E-F), which may be related to body size. There was a significant effect of genotype on time to fall in the ledge task (H_2_=12.505,p=0.001925), in which CD animals fell off the ledge faster than either WT (p=0.0071) or *Gtf2i\** (p=0.0069) littermates (Figure 3A). Similarly, there was a significant effect of genotype on the time spent balancing on a platform task (H_2_= 7.1578, p=0.02791) (Supplemental Figure 3G). Despite their comparable performance in strength and coordination tasks, the CD animals tended to have poorer balance, while the double mutants performed similar to WT animals. These findings suggest that other genes in the WSCR contribute to this balance deficit.

**Figure 3.**
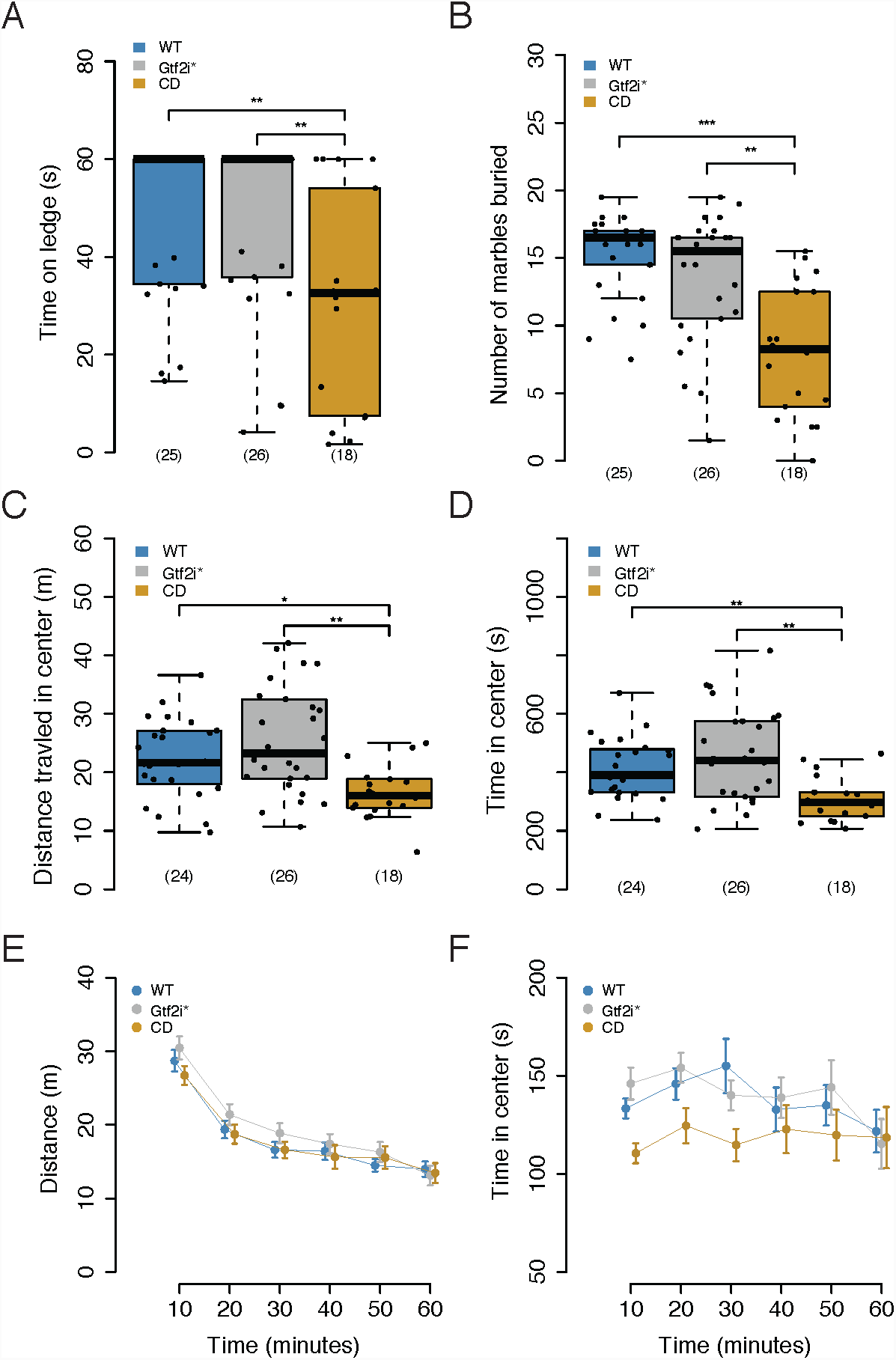
CD mice have motor deficits. **A** CD mice fall off a ledge sooner than WT or *Gtf2i\** mutants. **B** CD mice bury fewer marbles than either the WT or *Gtf2i\** mutants. **C** CD mice travel less distance in the center during marble burying task **D** CD animals spend less time in the center during marble burying task. **E** All genotypes travel similar distance in open field. **F** All genotypes spend similar time in the center during open field. * p < 0.05, ** p < 0.01, *** p < 0.001 Sample sizes are shown as numbers in parentheses

To test motor coordination in a more sensitive manner, we evaluated the mice on an accelerating rotarod. This task was performed over three days and tests coordination by quantifying how long a mouse can stay on a rotating rod. There was a main effect of day (F_2,339_ = 81.58, p< 2.2×10^−16^) and a main effect of sex (F_1,63_=10.0227, p = 0.002383), but no main effect of genotype (F_2,63_=2.0394, p=0.13861). We did not observe a sex by genotype interaction (F_2,63_=0.8155, p=0.447035) but did see a day by genotype interaction (F_4,333_=3.6270, p=0.006558). A post hoc comparison between genotypes within each day of testing showed that *Gtf2i*\* animals fell off more quickly compared to CD animals on day three (p=0.04) with no difference between WT and CD animals (Supplemental figure 3H). In contrast to the balance deficit seen on the ledge task but consistent with pole and screen performance, the rotarod results showed that all genotypes have similar motor coordination.

Marble burying is a species-specific behavior that assesses the natural tendency of mice to dig. This task also requires motor skills and has been used as a proxy for repetitive behaviors (45), which are seen in individuals with WS. It has been previously shown that CD animals bury fewer marbles than WT littermates (46, 47). Here we similarly show that there was significant effect of genotype in this task (F_2,66_=15.243, p=3.61×10^−6^). CD animals buried fewer marbles than both WT (p<0.001), and *Gtf2i\** mutants (p=0.000265) (Figure 3B), indicating that *Gtf2i*\* mutation is not sufficient to recapitulate CD phenotype. The differences in marble burying was not explained by any differences in activity levels between the genotypes during the task (F_2,65_=0.8974, p=0.4126; Supplemental Figure 3I). However, we did see a significant effect of genotype on distance traveled in the center of the apparatus (F_2,66_=13, p=0.0015), with CD mice traveling less distance in the center compared to WT (p=0.0301) and *Gtf2i\** (p=0.002) littermates (Figure 3C). There was also a corresponding significant effect of genotype on time spent in the center (F_2,66_=14.389, p=0.00075) with CD mice spending less time in the center than WT (p=0.0079) and *Gtf2i\** (p=0.0017) littermates. Avoidance of the center is generally interpreted in rodents as an increase in anxiety-like behavior (Figure 3D). Thus, these results provided further support to the hypothesis that genes besides *Gtf2i*\* contribute to an anxiety-related phenotype. It also suggested that the decreased marbles buried may be secondary to the decreased time in center and could reflect a phenotype secondary to anxiety rather than a direct stereotypy phenotype.

Finally, to test if the mutants have normal sensorimotor gating we looked at PPI. Similar to other tasks, contrasting evidence has been observed in WS mouse models in this task. Mouse of models of just *Gtf2i* showed no phenotype (33), whereas the proximal deletion mice showed decreased PPI; however, when combined with the distal deletion the phenotype that was suppressed (29). Here we show that all genotypes exhibited the expected increased PPI with an increasing pre-pulse stimulus (F_2,112_=620.61, p < 2e-16), but with no effect of genotype (F_2,56_=0.7742,p=0.466) or a pre-pulse by genotype interaction (F_4,112_=1.926,p=0.111) (Supplemental Figure J). A decrease was observed for overall startle response to the 120dB stimulus by CD animals, but when we included weight in the statistical model this effect disappeared (genotype F_2,55_=1.48, p=0.2365; weight F_1,55_=26,001, p=4.34e-6). Thus, the only phenotypic difference seen simply reflected the smaller size of the CD mice and not a change in sensorimotor gating (Supplemental Figure 3K).

### WSCR mutation does not produce robust anxiety-like behaviors

WS patients have heightened anxiety (42), and mouse models of *Gtf2i* (33, 35) and *Gtf2ird1* (30, 31) mutations have produced mixed evidence to support the role of these genes in anxiety phenotypes. Larger deletion models that have either the proximal or distal regions deleted showed anxiety-like phenotypes in the open field, but not in light-dark boxes (29). Similarly the CD model has been shown to not have any differences in the open field task (28). We wanted to directly compare animals with *Gtf2i* and *Gtf2ird1* mutations to CD animals in the same tasks to test exploratory and anxiety-like phenotypes. First, we looked at the behavior of the mice in an one hour locomotor activity task. We did not see any effect of genotype on the total distance traveled (F_2,66_=0.6324, p=0.53449), however there was a trend towards a time by genotype interaction (F_10,330_=1.7817, p=0.06283; Figure 3E) with the *Gtf2i\** mutants traveling further distance. This was consistent with the behavior observed during the three-chamber social approach task. In contrast to the marble burying task, here we did not see a significant main effect of genotype on the time spent in the center of the chamber (F_2,66_=2.3104, p=0.10720) though we observed a trend in the first ten minutes for CD mice to spend less time in the center (Figure 3F). However, the *Gtf2i\** mice did not show a similar trend. To further test for anxiety-like phenotypes, we performed elevated plus maze testing. Across the three days of testing, all genotypes spent similar percent time in the open arms of the apparatus (F_2,63_=0.6351, p=0.5332; Supplemental Figure 3L). Overall, our experiments indicate there may be a subtle increase on some tasks in anxiety-like behavior in CD mice. However, if there is such a phenotype, we see no evidence that *Gtf2i\** mutations are sufficient to produce it.

### GTF2I* mutation is not sufficient to reproduce WSCR mediated alterations of fear conditioning

Finally, as patients with WS have both intellectual disability and increased prevalence of phobias (42, 48), we tested associative learning and memory of the mice using a contextual and cued fear conditioning paradigm. These behaviors are also mediated by brain regions that have shown to be altered in mouse models of WS and human patients, namely the amygdala and hippocampus. Individuals with WS have altered structural and functional reactivity in the hippocampus and amygdala as reviewed in (2) compared to typically developing controls. Both of these regions play a role in both contextual and cued fear conditioning (49). Likewise, CD mice have been shown to have altered morphology and physiology in the hippocampus (28, 50), thought to be important in contextual fear conditioning.

We therefore tested associative learning and memory of the animals using a three day conditioned fear task (Figure 4A). During the conditioning trial on day one we saw a significant difference in baseline freezing during the first two minutes, when the mice were initially exploring the apparatus. There was a main effect of genotype (F_2,53_=5.31,p=0.00794) and a main effect of minute (F_1,53_=7.28, p=0.009), with the CD animals freezing more than the WT animals (p=0.04) and the *Gtf2i\** mutants (p=0.05) during minute one prior to any shock. By minute two of baseline, all animals showed similar levels of freezing. During the pairing of the foot shock with the context and tone during minutes three through five, we saw a significant effect of time (F_2,106_=100.3071, p < 2.2×10^−16^) and genotype (F_2,53_=3.4304, p=0.039723) as well as a time by genotype interaction (F_4,106_=3.9736, p = 0.004812). Specifically, all mice increased the amount of freezing after each foot shock, but after the last foot shock the *Gtf2i\** mutants froze less than the CD animals (p=0.002; Figure 4B), but similarly to the WT littermates. On the subsequent day, to test contextual fear memory, mice were put back in the same apparatus and freezing behavior was measured. Comparing the average of the first two minutes of freezing during fear memory recall on day two to the baseline of the conditioning day, we saw that all genotypes exhibited contextual fear memory; indicated by the increased levels of freezing when put back in the same context they were conditioned in (F_1,53_=36.4882, p=1.56×10^−7^; Supplemental Figure 4A). Looking across time during the fear memory recall we saw a significant effect of time (F_7,371_=2.7166, p=0.009291) with no main effect of genotype (F_2,53_=1.2507, p=0.294625), but a time by genotype interaction (F_14,371_=2.499, p=0.002085). Post hoc analysis within time showed that CD mice froze more than WT and *Gtf2i\** littermates during minute three of the task (Figure 4C).

**Figure 4.**
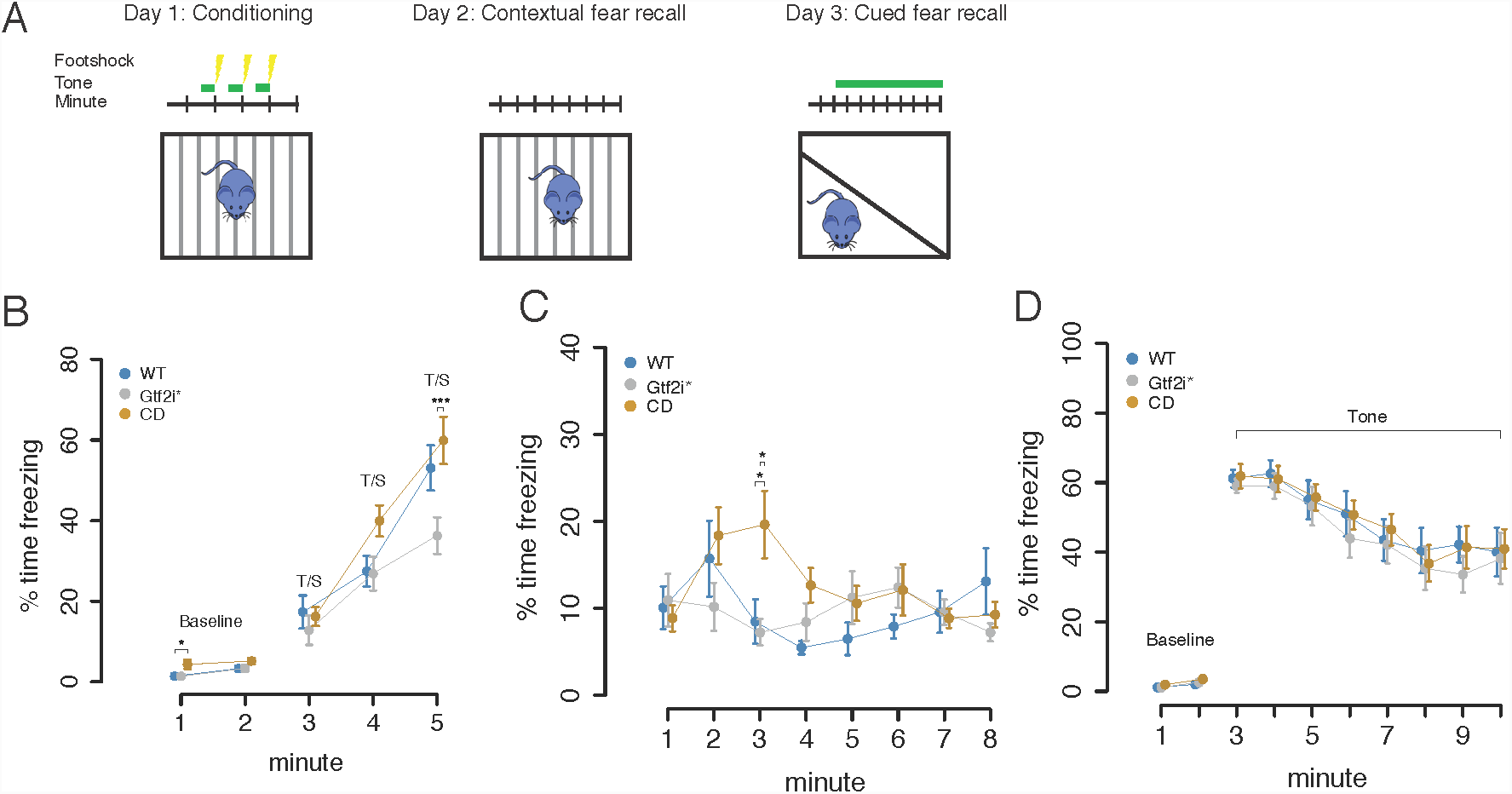
CD mice have more severe contextual fear phenotypes than double mutants. **A** The conditioned fear task design. Day one animals are delivered a tone and then a footshock throughout the five minute task. Day twp the animals are put in the same context without a footshock to measure contextual fear memory. Day three animals are put in a new chamber and delivered the tone to measure cued fear memory **B** Percent time freezing during conditioned fear acquisition. CD mice have increased baseline freezing during minute one and *Gtf2i\** mutants show decreased freezing during minute five **C** Percent time freezing during contextual fear memory recall. CD mice show elevated freezing during fear memory recall. **D** Percent time freezing during cued fear memory recall. All animals show increased freezing when the tone is played. * p < 0.05, ** p < 0.01, *** p < 0.001 Sample sizes are shown as numbers in parentheses

To test cued fear conditioning, on the subsequent day the mice were put in a different context and were played the tone that was paired with the foot shock during the conditioning day. All animals had similar freezing behavior during baseline (F_2,53_=1.061, p=0.353). For the duration of the tone, there was a significant effect of time (F_7,371_=21.5824, p<2×10^−16^) but no effect of genotype (F_2,53_=0.3014, p=0.741) or genotype by time interaction (F_14,371_=0.2128, p=0.999) (Figure 4D). Finally, the differences in freezing behavior could not be explained by sensitivity to the foot shock as all mice showed similar behavioral responses to increasing shock doses (F_2,56_=1.4521, p=0.2427; Supplemental Figure 4B). Overall, CD mice showed an enhancement of fear response to a contextual fear memory, and mutations in *Gtf2i*\* were not sufficient to reproduce this phenotype.

### Gtf2i* mutation is not sufficient to reproduce WSCR mediated alterations of hippocampal gene expression

In addition to permitting behavioral phenotyping, mouse models also allow for well powered and controlled examination of the molecular consequences of mutation in the environment of a fully developed and functioning central nervous system. Therefore, we turned from behavioral phenotyping of cognitive tasks to molecular phenotyping in the brains of these mice to 1) identify candidate molecular mediators of the behavioral phenotypes and 2) determine to what extent any transcriptional phenotype of WSCR mutation might be mediated by the haploinsufficiency of these two transcription factors. We specifically focused on the hippocampus, since we saw deficits in marble burying and differences in contextual fear memory, two behaviors thought to be mediated by hippocampal function (49, 51). Other studies in the CD animals have also shown there to be differences in LTP in the hippocampus as well as differences in Bdnf levels (47, 50). Yet the transcriptional consequences genome-wide of WSCR loss hav not been characterized in the hippocampus.

First, we conducted a targeted analysis of the genes in the WSCR locus. Of the 26 genes that make up the WSCR, only 15 were measurably expressed in the adult mouse hippocampus. Consistent with expectation, all genes in the WSCR region showed a decrease in RNA abundance in the CD animals, and genes that lie immediately outside the region were not affected. *Gtf2i*\* mutants only showed disruption of *Gtf2i* and *Gtf2ird1* in directions consistent with what was previously seen in our RT-qPCR. This confirmed the genotype of the samples, and indicated that these transcription factors are not robust trans regulators of any other genes in the locus (Figure 5A).

**Figure 5.**
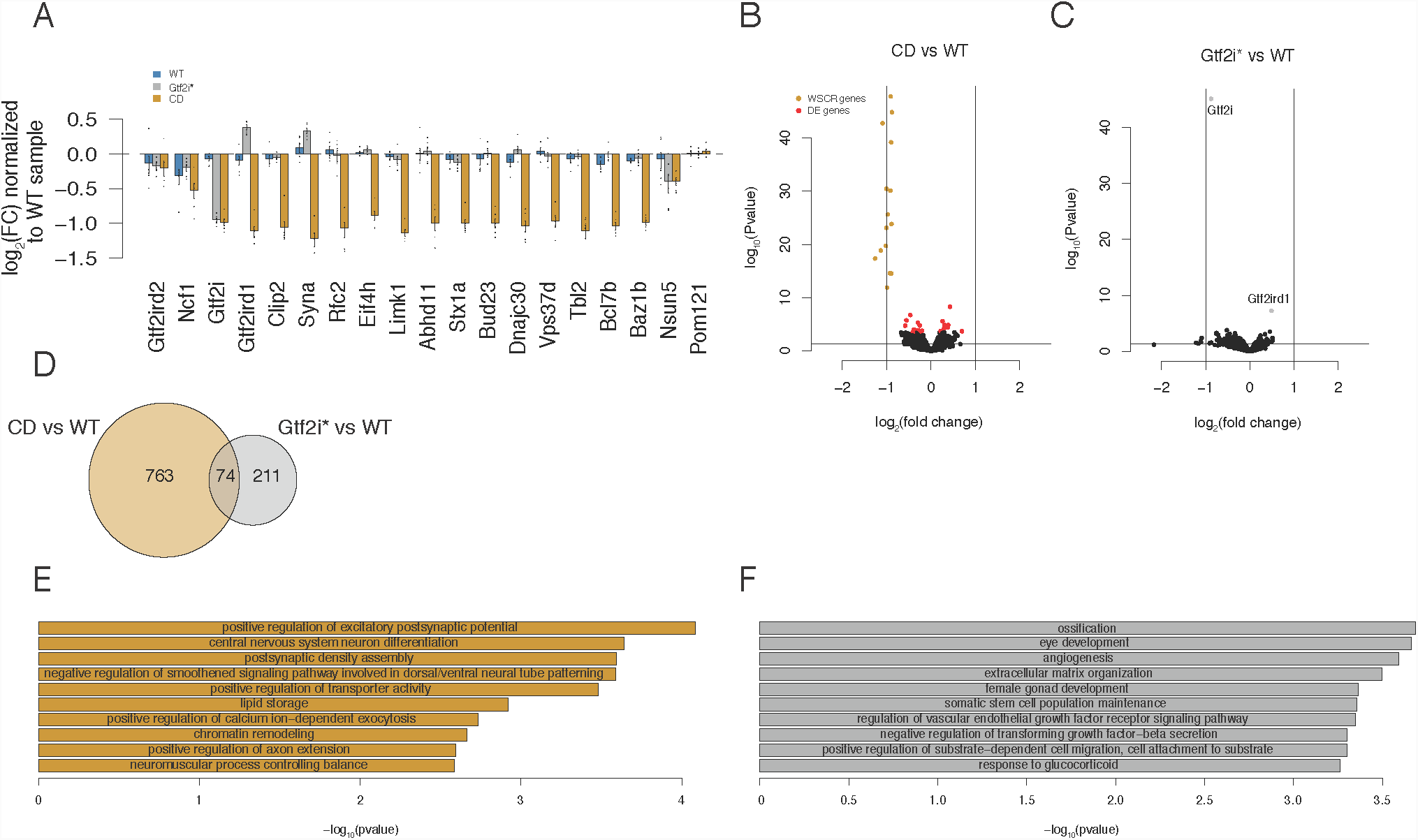
CD mice have altered mRNA for synaptic genes in a hippocampus transcriptome. **A** CD animals show decreased expression of the WSCR that are expressed in the hippocampus. **B** volcano plot comparing CD and WT differentially expressed genes.WSCR genes are highlighted in yellow and genes with FDR < 0.1 are highlighted in red. **C** Besides *Gtf2i* and *Gtf2ird1* there are no significantly differentially expressed genes **D** There is a 9% overlap between nominally significantly up and down regulated genes between CD and *Gtf2i*\* comparisons to WT controls. **E** CD differentially expressed genes are enriched for GO biological processes involved in synapses and nervous system development. **F** *Gtf2i\** differentially expressed genes are enriched for GO biological processed involved in more general organ development.

Next, we conducted differential expression analysis comparing WT to CD littermates to identify the molecular consequences of WSCR loss. At an FDR < 0.1 we found 39 genes to be differentially expressed. Of the 39 genes, 15 were genes that are located in the WSCR. This small number of differentially expressed genes was surprising given that several of the WSCR genes are described as transcription factors. In addition to these differentially expressed genes, the magnitude of the changes were small (Figure 5B and Supplemental Figure 5A). Interestingly, *Slc23a1* showed to be slightly but consistently more lowly expressed in the CD animals compared to the WT animals. This is a GABA transporter, suggesting that inhibitory signaling could be altered in the hippocampus. This gene has also been shown to decreased in WS-derived cortical neurons (23). Also of note, the *Iqgap2* gene was shown to be elevated in the CD animals compared to WT animals. This gene was also upregulated in WS iPSCs (24). We also looked at genes that have been investigated previously in the CD mouse, such as *Bdnf* and *Pi3kr* (46, 47) and we show that there was little change in gene expression between genotypes (Supplemental Figure 5B).

To determine if *Gtf2i*\* loss is sufficient to drive these transcriptional changes, we next examined differential expression comparing *Gtf2i*\* mutants to WT littermates. In contrast to WSCR mutation, we found only *Gtf2i* and *Gtf2ird1* to be differentially expressed at an FDR < 0.1 (Figure 5C). To get a broader idea of how similar the transcriptomes of the two genotypes are we compared the genes that are nominally up and downregulated between each mutant line and WT controls. We saw that there was about a 9% overlap between CD and *Gtf2i\** up and down regulated genes (Figure 5D). This is slightly below the amount of genes shown to be changed by *GTF2I* in iPSCs (24). Again this suggests that other genes in the WSCR are driving 90% of the transcriptional changes in the CD hippocampus.

To understand what role the nominally changed genes have in common we conducted a GO analysis. The biological processes that the CD genes were found to be involved in included synaptic functioning as well as nervous system differentiation. Interestingly processes that control balance were enriched and we and others have reported on balance deficits in CD animals (Figure 5E). When comparing these to 1000 random differential gene lists these biological processes are very specific to the genotype comparisons. For instance, out 1000 random test, positive regulation of excitatory synapses only occurred in the top 10 enriched GO terms two times (Supplemental Table 2). The cellular components that the genes are enriched for are extracellular, which is a similar result to the iPSC studies (24), as well as synapses. The molecular function ontologies which are enriched for the differentially expressed genes included calcium binding (Supplemental Figure 5). When comparing these to randomly determined gene expression changes, all but the extracelluar components seem to be specific to the CD versus WT comparison (Supplemental Table 2). In contrast, the *Gtf2i*\* GO analysis showed that these genes are enriched for more general organ system development and are not very nervous system specific (Figure 5F and Supplemental Table 3).

Overall, we have shown that the hemizygous loss of the WSCR has a mild but significant effect on the hippocampal transcriptome. Yet, the changes that do occur point to aberrations in synapses and nervous system development. Furthermore, loss of function mutations in *Gtf2i* and *Gtf2ird1* have an even smaller effect on the transcriptome and can only account for 9% of the changes incurred by loss of the WSCR.

## Discussion

Contiguous gene disorders such as WS provide insight into regions of the genome that have large effects on specific aspects of human cognition and behavior. The specific cognitive profile of WS is characterized by deficits in visual-spatial processing with relative strengths in language, and the archetypal behavioral profile consists of increased social interest, strong eye contact, high levels of anxiety, and in some cases specific phobias and hyperactivity. Here we used a new mouse model to test if loss of the paralogous transcription factors *Gtf2i* and *Gtf2ird1* are sufficient to phenocopy the behaviors and transcriptomic changes of mice that lack the entire WSCR.

Overall, CD mice consistently have more severe phenotypes than the *Gtf2i\** mutants. We saw that the CD animals have a deficit in social communication as measured by maternal separation induced pup ultrasonic vocalizations. The *Gtf2i\** mutants on average make fewer calls than the WT littermates, however not significantly so, but this may suggest that these two transcription factors contribute slightly to this phenotype but other genes in the region are necessary to produce the full phenotype seen in the CD animals. Previously it was shown that animals that have increased copy number of *Gtf2i* increased the number of pup USVs emitted while animals with only one copy produced similar number of calls to WT animals (34). This was interpreted as increased separation anxiety. Here we see that lower copy number of the entire region produces the opposite effect of increased *Gtf2i* copy number. Decreased USVs could mean there is a lack of motivation to make the calls or an inability to make as many calls. A model of *Gtf2ird1* mutant animals was shown to have different USV production due to a difference in the muscle composition of the larynx (32). This has not been shown in the CD animals but it could contribute to the phenotype as well as differences in the skull morphology (28). Another possible explanation is that since the production of USVs is a developmentally regulated trait, it could be that deleting 26 genes could disrupt typical developmental trajectories. While we do not see any gross developmental problems such as lower weight or delayed detachment of pinnae, the deletion could have a more severe effect on brain development, thus affecting developmentally regulated behavioral traits.

To our surprise, there was no detectable social phenotype in the *Gtf2i\** mutants or CD animals in the classical three-chamber social approach assay. Our results showed that all genotypes on average prefer to investigate the social stimulus for a similar amount of time. The preference for social novelty is also intact across all the groups. In an attempt to test if the WS models fail to habituate to a social stimulus we showed that after thirty minutes of having the opportunity to investigate an unfamiliar mouse or an empty cup, all genotypes habituate to the social stimulus and by the end of the thirty minutes seem to have a small preference for the empty cup. The three-chamber social approach task has been done in the larger partial deletion models where they have shown that the proximal deletion and the trans full deletion models have a significant preference for the social stimulus, and the WT and distal deletion mice do not show a preference, suggesting that the proximal deletion, which harbors genes such as *Gtf2i* and *Gtf2ird1*, are involved in this social task (29). Mouse models that are haploinsufficient for only *Gtf2i* have shown in the three-chamber approach task that after eight minutes WT animals investigate a novel object the same amount as a social stimulus, but the *Gtf2i* mutants still have a significant preference suggesting a lack of habituation (33). In another *Gtf2i* model, Martin et al. compared animals with one, two, three, and four copies of *Gtf2i* in the three-chamber social approach task, and showed that only animals with one and three copies of *Gtf2i* displayed a significant preference for the social stimulus (36), but WT animals did not. These three-chamber social approach tests are interpreting a lack of significance as evidence for increased social behavior and not directly comparing the levels of investigation between genotypes (52). Furthermore, in some cases the WT controls are not showing the expected preference for the social stimulus, thus, possibly confounding interpretation of the mutant preference.

The three-chamber social approach assay has come under recent criticism due to how dependent it is on activity levels of mice and its lower heritability compared to tests of direct social interaction (53). The CD animals had not previously been tested in this procedure excatly, but have been tested in a modified social approach where the time spent investigating a mouse in a cup is measured but with no competing non-social stimulus (28, 46, 47). The data showed that the CD animals investigated the social stimulus for more time than the WT animals and delivery of *Gtf2i* cDNA by AAV9 via the magna cisterna can return the investigation time to normal levels (46). Here, we showed that all animals preferred the social stimulus. It is possible that the standard social approach suffers from several confounding factors, such as lower heritability, as well as activity and anxiety-like components that make this task less sensitive to detect a hypersocial phenotype in WS models. It could also be that the three-chamber social task does not test the specific aspects of social behavior that are disrupted in WS models. For example, newer tasks, such as social operant tasks that test motivation to receive a social stimulus may more directly test the aspects of social behavior that are affected in WS. This task has been performed on *Gtf2i* mutants and mice that have only one copy of *Gtf2i* will work harder to receive a social reward (36).

Direct social tasks have higher heritability than the three-chamber social approach and offer a more natural social experience (53), which may make them a more sensitive assay for testing social behaviors. Direct tasks have shown that *Gtf2i* models have increased nose-to-nose investigation time (36), mouse models lacking the proximal end of the region have increased investigation frequency (29), and *Gtf2ird1* mutants make fewer aggressive actions but show increased following time (30). We employed the resident-intruder paradigm as a full contact social assay. While we did not see bouts of aggression from any of the genotypes, we could see differences in social investigation. To our surprise, the CD animals spent less time overall in anogenital sniffing and nose-to-nose sniffing of the intruder animals when compared to WT littermates. The double mutants were not significantly different from the WT animals but had intermediate values between the WT and CD animals. This phenotype was being driven by the decreased time per bout of investigation in the CD animals, as all genotypes had a similar frequency of the sniffing behavior. This result was contrary to what would be predicted from the human condition and previous mouse results. However, while individuals with WS are described as having prosocial behavior in terms of increased social approach and friendliness (54), they also have difficulties maintaining long term relationships because of deficits in other aspects of social behavior (4, 5, 55, 56), and on scales measuring social reciprocity often score in the autistic range (5). In addition, there is a high co-morbidity with ADHD which has features of impulsiveness (57). While the CD animals did not show the expected increase in social interest, this may be a manifestation of attention deficits that are present from deleting the 26 genes in the WSCR, but this needs to be examined. Loss-of-function mutations in *Gtf2i* and *Gtf2ird1* were not sufficient to produce as strong an effect in these investigative behaviors. However, the somewhat intermediate effect suggests they could contribute to it.

One limitation of our study is that some aspects of the social phenotype in the models tested here could be masked by the mouse background strain. While we have controlled for mouse background strain in our experiments by only using the F1 generation of the FVB/AntJ and C57BL/6J cross, the hybrid background may prevent the manifestation of a social phenotype caused by the mutations tested. For example, it has been documented that craniofacial phenotypes in *Gtf2ird1* models are sensitive to background strain (13, 30, 41, 44). Here, the double mutants and CD animals on the hybrid background showed no dominance phenotype in the tube test. However, when we tested each mutation on the respective mouse background strain, we saw that the CD animals had a submissive phenotype, but the double mutants did not. Studies done in the larger partial deletions have shown altered win/loss ratios in the tube test in the proximal deletion and full trans deletion models (29), suggesting that the CD models on the C57BL/6J background can replicate this phenotype, but other genes in the proximal region besides *Gtf2i* and *Gtf2ird1* are also required.

In this study, we have replicated several of the phenotypes previously seen in the CD animals, such as marble burying and balance deficits (28, 47, 50). It was shown that CD animals bury fewer marbles than WT animals and rescuing the *Gtf2i* levels in the hippocampus did not rescue this phenotype. Both the results presented here and in Borralleras et al. suggest that other genes in the region beyond *Gtf2i* and *Gtf2ird1* are important in this behavior. Here we have extended the results to suggest that there could be an anxiety-like component to the marble burying deficit. By tracking the animals during the task we see that CD animals spend less time and travel less distance in the center of the apparatus. This could preclude them from burying as many marbles in the center. It could also be that the CD animals do not show the normal motivation to dig.

CD animals showed difficulty in balancing tasks, but normal motor coordination. Motor coordination of WS has been tested using the rotarod. The larger partial deletion models showed that the distal deletion and proximal deletion mice had intermediate phenotypes with the full trans deletion mice falling off the rotarod sooner (29). Similarly the CD mice have shown deficits in the rotarod and addition of *Gtf2i* coding sequence does not rescue this phenotype (50). The CD mice in this study did not show a deficit in the rotarod despite having poor balance on the ledge and platform tasks. CD animals were not able to balance on a ledge or platform as long as their WT and *Gtf2i\** mutant littermates. This suggests that motor coordination, as tested by our rotarod paradigm, is intact in these WS models, but balance is specifically affected in the CD animals. The discrepancy could be due to body size. The adult CD animals are significantly smaller than the WT and *Gtf2i\** mutants, which could make staying on the wider rotarod less challenging. This study also used a different accelerating paradigm where the rod itself is continuously accelerating until the mouse falls off while other paradigms test the mice at different continuous rotation speeds.

Along with balance and coordination problems, individuals with WS tend to have specific phobias and high levels of non-social anxiety (42). We showed that CD animals had an altered fear conditioning response. We saw that the CD animals have an increased fear response in contextual fear but not cued fear. It was previously reported that CD animals showed a slight decrease in freezing but was not significant (28). Two separate *Gtf2ird1* mutations have shown contrasting results, one showed an increased fear response (16) while another showed decreased fear response (30). It could be that this hybrid background used here is more sensitive to see increases in freezing because FVB/AntJ do not exhibit as much freezing in conditioned fear tasks as C57BL/6J animals (58). The observed increased contextual fear response could be due to differences in the hippocampus and amygdala, both regions that have been shown to be disrupted in WS. We did not see a robust anxiety like behavior phenotypes in one hour locomotor task or the elevated plus maze, which is consistent with previous findings in the CD model (28). However, we did see reduced time and distance traveled in the center during the marble burying task. Perhaps suggesting that the novel environment in combination with the novel marbles can induce slightly higher levels of anxiety in the CD model.

Given the behavioral differences in marble burying and contextual fear, two behaviors thought to be mediated by the hippocampus (49, 51), we examined the transcriptomes of the hippocampus of the *Gtf2i\** mutants and CD animals and compared them to WT littermates. This provided the first transcriptional profile documenting the consequences of the 26 gene deletion in a mature brain, and allowed us to determine what portion of that was driven by Gtf2i* proteins. Surprisingly, we did not see any significantly differentially expressed genes between the *Gtf2i\** mutants and WT littermates, besides the mutated genes themselves. Looking at the overlap of nominally differentially expressed genes between CD-WT and *Gtf2i*-*WT comparisons, showed a small overlap of about 9%. This is slightly less than the estimate from Adamo *et al.*, of 15-20% of genes dysregulated in WS iPSCs being attributed to reduced levels of *GTF2I*. Perhaps these general findings suggest that *Gtf2i* and *Gtf2ird1* contribute to small transcriptional changes broadly across the genome, and in combination with other genes in the WSCR more profound neural specific gene disruptions become apparent.

Our transcriptional studies overall showed limited impact of *Gtf2i*\* mutation in the brain. The global brain transcriptome of *Gtf2i* mutants has not been investigated, but brain transcriptome studies of *Gtf2ird1* knockout mouse models have not found any evidence of differentially expressed genes (59). These data suggest that in the adult hippocampus these two transcription factors do not greatly affect the transcriptome. There are some limitations to this negative result. It could be that we are diluting some of the signal because we are studying the effects on the transcriptome of the whole hippocampus, which has a diverse cellular composition. Larger effect sizes might be detected in more homogenous cellular populations. Likewise, if these genes regulate dynamics of gene expression rather than baseline values, greater differences might become apparent after experimental manipulations that activate transcription.

One additional limitation of our study is that the mutated *Gtf2ird1* allele is still producing an N-terminally truncated protein. However, we show that N-truncated Gtf2ird1 does not bind to its known target, the promoter region of *Gtf2ird1*, and this absence leads to increased RNA from the locus, consistent with a loss of its transcriptional repressor function. Thus, we confirmed this truncated protein is a loss of function for the only known roles for Gtf2ird1. However, it is possible that the protein does have other unknown functions we could not assay here. It has also been proven to be a remarkably challenging gene to completely disrupt, across multiple studies (30, 38). The combination of the upregulation of its RNA upon deletion with the ability to re-initiate at a variety of downstream codons is intriguing. One possibility is that Gtf2ird1 has an unusual amount of homeostatic regulation at both transcriptional and translational levels that are attempting to normalize protein levels. Another possibility is that these kinds of events are actually quite common across genes, but that they are detected in Gtf2ird1 because the WT protein is at such low abundance it is on par with what is actually an infrequent translation reinitiation event. Our detection of Gtf2ird1 protein in the brain required substantial optimization and is still only apparent in younger brains. Indeed, in validations of mutations of more abundant proteins, the immunoblots may not be routinely developed long enough to see a trace re-initiation event that might occur. Regardless, future studies aimed at understanding the transcriptional and translational regulation of this unusual gene would be of interest.

Examining the profile of CD mutants compared to WT littermates, we do define a number of transcriptionally dysregulated genes. Of the genes in WSCR that are expressed in the hippocampus all had decreased expression in the CD animals. In addition, there were 24 genes outside the WSCR that had a FDR < 0.1 between CD and WT controls. Among these genes is *Slc23a1*, the GABA vesicle transporter, which is down regulated in CD animals. Interestingly this gene was also found to be down regulated in human iPSC derived neurons from individuals with WS (23). This points to aberrant inhibitory activity in the CD brain, which could lead to functional deficits. Also consistent with other human WS derived iPSC studies, the gene *Iqgap2* was shown to be upregulated in the CD hippocampus (24), and has the potential to interact with the cytoskeleton through actin binding (60). Broadening the analysis to include nominally differentially expressed genes and conducting systems-level analyses, the CD-WT comparison highlighted genes involved in the positive regulation of excitatory postsynaptic potential. Chailangkarn et al. showed that WS derived iPSC neurons had increased glutamatergic synapses. Our data also showed some signal in the GO term for postsynaptic density assembly. Taken together these data suggest abnormal synapse functioning in the CD animals and potentially altered inhibitory/excitatory balance. This also suggests pharmacological agents that increase GABA tone may be of use in reversing some WS phenotypes. The RNA-seq data also had signal in neuromuscular processes controlling balance. Altered gene expression in the CD animals could be contributing to the balance deficits. In contrast to the synapse and neural specific GO term enrichment seen in the CD-WT comparison, comparing the transcriptomes of the *Gtf2i\** mutants and WT shows signal in more general organ development, such as ossification and eye development.

Taken together, our results support the hypothesis that other genes in the WSCR besides *Gtf2i* and *Gtf2ird1* are necessary to produce some phenotypes that are seen when the entire WSCR is deleted. While these two transcription factors have been highlighted in the human literature as large contributors to the WS phenotype, the literature is also consistent with a model where most genes contribute to aspects of different phenotypes in WS, but the full phenotypic effects occur when all the genes are deleted (Figure 6). Studying patients with atypical deletions highlights the variability of the region. Even within families that have inherited small deletions some of the cardiovascular, cognitive, and craniofacial phenotypes have incomplete penetrance (12, 14, 19). Comparing the deletion sizes and corresponding phenotypes shows a large overlap of genes that are deleted, but no clear pattern of which specific phenotypes are affected. Many of atypical deletions described to date that do not have *Gtf2i* and *Gtf2ird1* deleted show no overfriendly phenotype, but there are examples where this is not true. Recent work in zebrafish that was done to dissect which genes in the 16p11.2 region contribute to craniofacial dysmorphology led to a similar conclusion, that multiple genes in the region contribute to the phenotype but in combination some have synergistic effects and others have additive effects (61). Sanders *et al.* also suggested that copy number variations with higher gene content are more likely to have several genes of smaller effect sizes suggesting an oligogenic pattern of contribution (7). Our data suggests that looking beyond the general transcription factor 2I family at possible combinations of more genes in the region may more completely reproduce the WS phenotype. Given the ease of making new mouse models with current genome editing technology, a combinatorial dissection of the region is feasible and could lead to interesting new insight into the underlying mechanisms that contribute to the phenotypic spectrum of WS.

**Figure 6.**
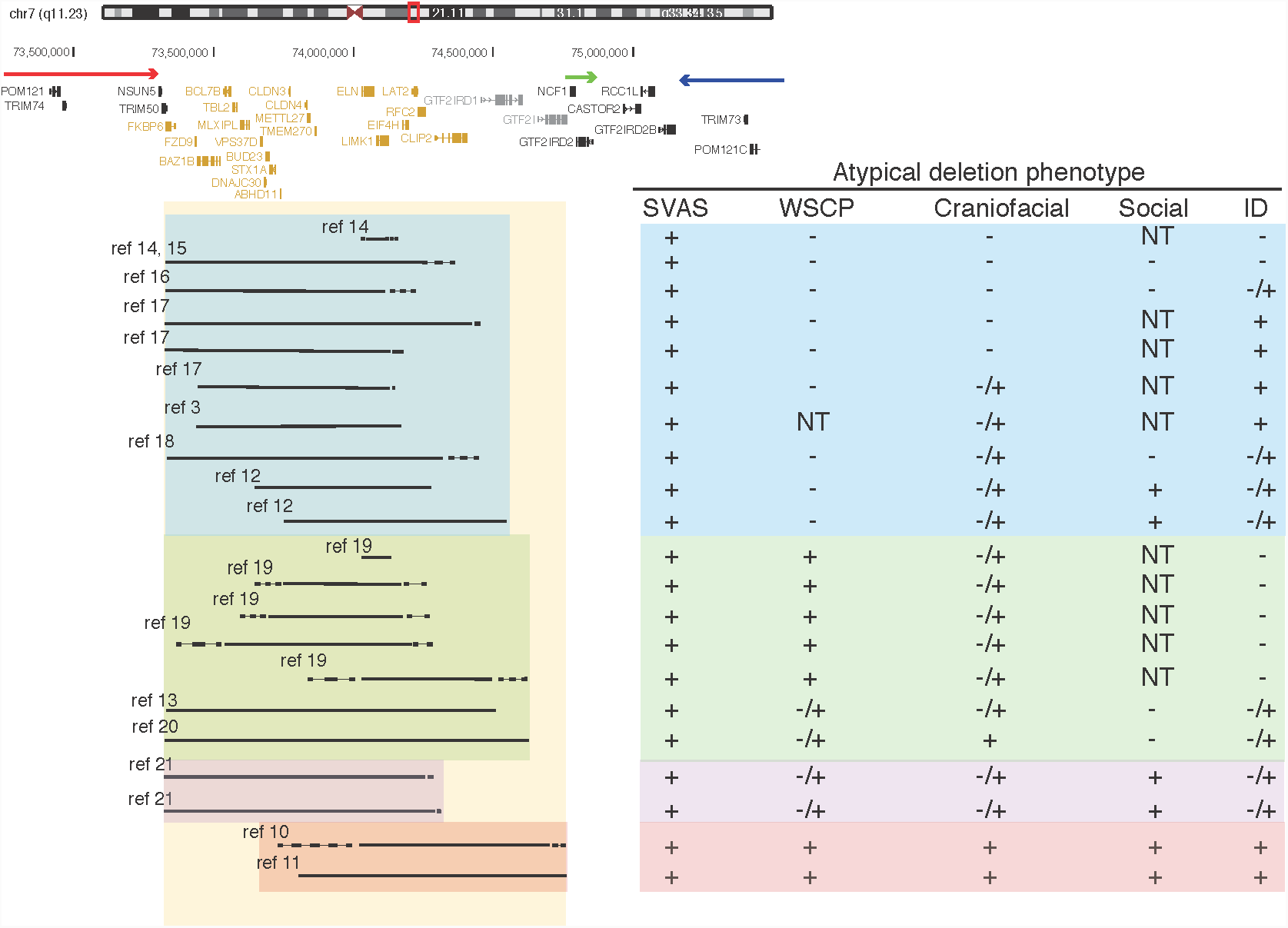
Human atypical deletions support oligogenic contribution of genes in the WSCR to phenotypes. Schematic of the WSCR on chr7q11.23. The arrows indicate the regions of low copy repeats. The typical deletion is demarcated using the yellow box. Atypical deletions demarcated in blue show no contribution to the WSCP. Atypical deletions demarcated in green show contribution to the WSCP. Atypical deletions demarcated in purple provide evidence of deletions that spare *GTF2I* and *GTF2IRD1* that show contributions to across phenotypic domains including social behavior. Atypical deletions demarcated in red provide evidence that the telomeric region is sufficient to produce the full spectrum of phenotypes. The large amount of overlap of all deleted regions and the mild phenotypes present across the atypical deletions suggests an oligogenic pattern. SVAS (supravalvular aortic stenosis), WSCP (Williams syndrome cognitive pfofile) ID (intellectual disability) NT (Not tested), - absent, + present, -/+ milder than typical WS.

## Materials and Methods

### Generating genome edited mice

sgRNAs were designed to target early constitutive exons of the mouse *Gtf2i* and *Gtf2ird1* genes. The gRNAs were cloned into the pX330 Cas9 expression plasmid (Addgene) and transfected into N2a cells to validate the cutting ability of each gRNA using the T7 enzyme assay. Primers used to amplify target regions tested by the T7 enzyme assay are in Supplemental Table 4. One guide was selected for each gene based on cutting activity (Supplemental Table 4). The gRNAs were in vitro transcribed using MEGAShortScript (Ambion) and Cas9 mRNA was in vitro transcribed, G-capped, and poly-A tailed using the mMessageMachine kit (Ambion). The mouse genetics core at Washington University School of Medicine co-injected the Cas9 mRNA (25ng/ul) along with both gRNAs (13ng/ul of each gRNA) into FVB/NJ fertilized eggs and implanted the embryos into recipient mothers. This resulted in 57 founders. Founders were initially checked for any editing events using the T7 assay. There were 36 animals with no editing events. We deep sequenced the expected cut sites, as described below, in the remaining 21 founders to identify which alleles were present.. Founders were crossed to wild type (WT) FVB/AntJ (https://www.jax.org/strain/004828) animals, which are different from FVB/NJs at two loci; Tyr^c-ch^ results in a chinchilla coat color and they are homozygous WT for the 129P2/OlaHSd *Pde6b* allele, which prevents them from developing blindness due to retinal degeneration. Coat color was visually genotyped and the functional FVB/AntJ *Pde6b* allele was genotyped using primers recommended by Jackson labs (Supplemental Table 5). The mice were crossed to FVB/AntJ until the mutations were on a background homozygous for the FVB/AntJ coat color and *Pde6b* alleles.

### Genotyping

Initial founder genotyping was performed by deep sequencing amplicons around the expected cuts sites of each gRNA. Primers were designed around the cut sites using the NCBI primer blast tool. To allow for Illumina sequencing we concatenated the Illumina adapter sequences to the designed primers (Supplemental Table 5). The regions surrounding the cut sites were amplified using the following thermocycler conditions: 95° C 4 minutes, 95° C 35 seconds, 58.9° C 45 seconds, 72° C 1 minute 15 seconds, repeat steps 2 through 4 35 times, 72° C for 7 minutes, hold at 4° C. A subsequent round of PCR was performed to add the requisite Illumina P5 and P7 sequences as well as sample specific indexes using the following thermocycler conditions: 98° C 3 minutes, 98° C 10 seconds, 64° C 30 seconds, 72° C 1 minute, repeat steps 2 through 4 20 times, 72° C 5 minutes, hold 4° C. The PCR amplicons were pooled and run on a 2% agarose gel and the expected band size was gel extracted using the NucleoSpin gel extraction kit (Macherye-Nagel). The samples were sequenced on a MiSeq. The raw fastq files were aligned to the mm10 genome using bwa v0.7.17 –mem with default settings (62), and the bam files were visualized using the integrated genome visualizer (IGV)v2.3.29 to determine the genotype.

Once the alleles of the founder lines were shown to be in the germline, we designed PCR genotyping assays that can distinguish mutant and WT alleles. Since the *Gtf2i* mutation and the *Gtf2ird1* mutation are in linkage and are always passed on together, primers were designed that would only amplify the five base pair deletion in exon three of *Gtf2ird1*. The primer was designed so that the three prime end of the forward primer sits on the new junction formed by the mutation with an expected size of 500bp. Beta actin primers, with an expected size of 138bp, were also used to help ensure specificity of the mutation specific *Gtf2ird1* primers as well as act as a PCR control (Supplemental Table 5). The CD animals were genotyped using primer sequences provided by Dr. Victoria Campuzano and primers that amplify the WT *Gtf2ird1* allele as a PCR control (Supplemental Table 5).

PCR was performed on toe clippings that were incubated overnight at 55° C in tail lysis buffer (10mM Tris pH 8, 0.4M NaCl, 2mM EDTA, 0.1% SDS, 3.6U/mL Proteinase K (NEB)). The proteinase K was inactivated by incubation at 99° C for 10 minutes. 1ul of lysate was used in the PCR reactions. Two bands indicated a heterozygous mutation in *Gtf2i* and *Gtf2ird1*. The cycling conditions for the 5bp *Gtf2ird1* deletion were: 95° C 4 minutes, 95° C 35 seconds, 66.1° C 45 seconds, 72° C 1 minute 15 seconds, repeat steps 2 through 4 35 times, 72° C for 7 minutes, hold at 4° C. The cycling conditions for the CD genotyping were: 95° C 4 minutes, 95° C 35 seconds, 58° C 45 seconds, 72° C 1 minute 15 seconds, repeat steps 2 through 4 35 times, 72° C for 7 minutes, hold at 4° C.

### Western blotting

E13.5 whole brains were dissected in cold PBS and immediately frozen in liquid nitrogen and stored at −80°C until genotyping was performed. Frozen brains were homogenized in 500ul of 1x RIPA buffer (10mM Tris HCl pH 7.5, 140mM NaCl, 1mM EDTA, 1% Triton X-100, 0.1% DOC, 0.1% SDS, 10mM Na_3_V0_4_, 10mM NaF, 1x protease inhibitor (Roche)) and RNAase inhibitors (RNasin (Promega) and SUPERase In (Thermo Fisher Scientific) and incubated on ice for 20 minutes. Lysates were cleared by centrifugation at 10,000g for 10 minutes at 4° C. The lysate was split into two 100ul aliquots for protein analysis and 250ul of lysate was added to 750ul of Tizol LS (Thermo Fisher Scientific) for RNA analysis. Protein concentration was quantified using a BCA assay and loaded at 25- 50ug in 1x Lamelli Buffer with B-mercaptoethanol onto a 4-15% TGX protean gel (Bio-Rad). In some experiments to achieve greater separation to detect the N-truncation, the protein lysates were instead run on a 7.5% TGX protean gel (Bio-Rad). The protein was transferred to PVDF 0.2um membrane by wet transfer. The membrane was blocked for one hour at RT with TBST 5% milk. The membranes were cut at 75KDa, and the top of the membrane was probed for either Gtf2i or Gtf2ird1, and the bottom of the membrane was probed for Gapdh, with the following primary antibodies: Rabbit anti-GTF2IRD1 (1:500, Novus, NBP1-91973), Mouse anti-GTF2I (1:1000 BD Transduction Laboratories, BAP-135), and Mouse anti-Gapdh (1:10,000, Sigma Aldrich, G8795). Primary antibodies were incubated overnight at 4° C in TBST 5% milk. We used the following secondary antibodies: HRP-conjugated Goat anti Rabbit IgG (1:2000, Sigma Aldrich, AP307P) and HRP-conjugated Goat anti Mouse IgG (1:2000, Bio Rad, 1706516) and incubated for 1 hour at room temperature. Signal was detected using Clarity Western ECL substrate (Bio-Rad) in a MyECL Imager (Thermo Scientific). Quantification of bands was performed using Fiji (NIH) (63) normalizing to Gapdh levels and a WT reference sample.

### Transcript measurement using RT-qPCR

Total RNA from E13.5 brains lysates was extracted from Trizol LS using the Zymo Clean and Concentrator-5 with on column DNAase I digestion and eluted in 30ul of water. RNA quantity and purity was determined using a Nanodrop 2000 (Thermo Scientific). cDNA was prepared using 1ug of total RNA and the qscript cDNA synthesis kit (Quanta Biosciences). 25ng of cDNA was used in a 10ul RT-qPCR reaction with 2x PowerUP SYBR Green Master Mix (Applied Biosystems) and 500nM primers that would amplify constitutive exons of *Gtf2ird1* (exons 8/9), *Gtf2i* (exons 25/27) or *Gapdh* (Supplemental Table 5). The RT-qPCR was carried out in a QuantStudio6Flex machine (Applied Biosystems) with the following cycling conditions: 95° C 20 seconds, 95° C 1 second, 60° C 20 seconds, repeat steps 2 through 3 40 times. There were three biological replicates per genotype in all experiments and each cDNA was assessed in triplicate technical replicates. Relative transcript abundance of *Gtf2i* and *Gtf2ird1* was determined using the deltaCT method normalizing to *Gapdh*.

### ChIP-qPCR

#### Chromatin preparation

Chromatin was prepared by homogenizing one frozen E13.5 brain in 10mL of 1x cross-linking buffer (10mM HEPES pH7.5, 100mM NaCl, 1mM EDTA, 1mM EGTA, 1% Formaldehyde (Sigma)) using the large clearance pestle in a Dounce homogenizer and allowed to crosslink for 10 minutes at room temperature with end-over-end rotation. The formaldehyde was quenched with 625ul of 2M glycine. The cells were spun down at 200g at 4° C and the pellet was washed with 10mL 1x PBS 0.2mM PMSF and spun again. The pellet was resuspended in 5mL L1 buffer (50mM HEPES pH 7.5, 140 mM NaCl, 1mM EDTA, 1mM EGTA, 0.25% Triton X-100, 0.5% NP40, 10.0% glycerol,1mM BGP (Sigma), 1x Na Butyrate (Millipore), 20mM NaF, 1x protease inhibitor (Roche)) and homogenized using the low clearance pestle in a Dounce homogenizer to lyse the cells and leave the nuclei intact. The homogenate was spun at 800g for 10 minutes at 4° C to pellet the nuclei. The pellet was washed in 5mL of L1 buffer and spun again and resuspended in 5mL of L2 buffer (10mM Tris-HCl pH 8.0, 200mM NaCl, 1mM BGP, 1x Na Butyrate, 20mM NaF, 1x protease inhibitor) and incubated at room temperature for 10 minutes while shaking. The nuclei were pelleted by spinning at 800g for 10 minutes and resuspended in 950ul of L3 buffer (10mM Tris-HCl pH 8.0, 1mM EDTA, 1mM EGTA, 0.3% SDS, 1mM BGP, 1x Na Butyrate, 20mM NaF, 1x protease inhibitor) and transferred to a milliTUBE 1mL AFA Fiber (100)(Covaris). The sample was then sonicated to a DNA size range of 100-500bp in a Covaris E220 focused ultrasonicator with 5% duty factor, 140 PIP, and 200cbp. The sonicated samples were diluted to 0.1% SDS using 950ul of L3 buffer and 950ul of 3x Covaris buffer (20mM Tris-HCl pH 8.0, 3.0% Triton X-100, 450mM NaCl, 3mM EDTA). The samples were spun at max speed in a tabletop centrifuge for 10 minutes at 4° C to pellet any insoluble matter. The supernatant was pre-cleared by incubating with 15ul of protein G coated streptavidin beads (ThermoFisher) for two hours at 4° C.

#### Chromatin IP

GTF2IRD1 antibody (Rb anti GTF2IRD1 NBP1-91973 LOT:R40410) was conjugated to protein G coated streptavidin beads by incubating 6ug of antibody (10ul) with 15ul of beads in 500ul TBSTBp (1x TBS, 0.1% Tween 20, 1%BSA, .2mM PMSF) and end-over-end rotation for one hour at room temperature. The antibody-conjugated beads were washed three times with 500ul of TBSTBp. 400ul of the pre-cleared lysate was added to the antibody-conjugated beads and rotated end-over-end at 4° C overnight. 80ul of the precleared lysate was added to 120ul of 1x TE buffer with 1% SDS and frozen overnight to be the input sample.

The IP was washed two times with a low salt buffer (10mM Tris-HCl pH 8.0, 2mM EDTA, 150mM NaCl, 1.0% Triton X-100, 0.1% SDS), two times with a high salt buffer (10mM Trish-HCl pH 8.0, 2mM EDTA, 500mM NaCl, 1.0% Triton X-100, 0.1% SDS), two times with LiCl wash buffer (10mM Tris-HCl pH 8.0, 1mM EDTA, 250mM LiCl (Sigma), 0.5% NaDeoxycholate, 1.0% NP40), and one time with TE (10mM Tris-HCl pH 8.0, 1mM EDTA) buffer. The samples were eluted from the beads by incubating with 100ul of 1x TE and 1% SDS in an Eppendorf thermomixer R at 65° C for 30 minutes, mixing at 1400rpm. This was repeated for a total of 200ul of eluate. The samples and input were then de-crosslinked by incubating in a thermocycler (T1000 Bio-Rad) for 16 hours at 65° C. The samples were incubated with 10ug of RNAseA (Invitrogen) at 37° C for 30 minutes. The samples were then incubated with 140ug of Proteinase K (NEB) at 55° C in a thermomixer mixing at 900rpm for two hours. The DNA was extracted using phenol/chloroform/isoamyl alcohol (Ambion) and cleaned up using Qiagen PCR purification kit and eluted two times using 30ul of buffer EB for a total of 60ul. The concentration was assessed using the highsensitivity DNA kit for qubit (Thermo Fisher Scientific). A portion of the input DNA was run on a 2% agarose gel post stained with ethidium bromide to check the DNA fragmentation.

#### ChIP qPCR

Primers were designed to amplify the region around the *Gtf2ird1* transcription start site (TSS), which has been shown to be a target of Gtf2ird1 binding (38). Two primer sets were also designed to amplify off target regions, one 10kb upstream of the *Bdnf* TSS and one 7Kbp upstream of the *Pcbp3* TSS. These were far enough away from any TSS that it would be unlikely that there would be a promoter region. The primers can be found in Supplemental Table 5. A standard curve was made by diluting the input sample for each IP sample 1:3, 1:30, and 1:300. The input, the input dilutions, and the IP samples for each genotype condition were run in triplicate using the Sybr green Power UP mastermix (AppliedBiosystems) and primers at a final concentration of 250nM. The PCR was carried out in a QuantStudio6Flex machine (Applied Biosystems) with the following cycling conditions: 50° C for 2 minutes, 95° C for 10 minutes, 95° C 15 seconds, 60° C for 1 minute, repeat steps 3 through 4 40 times. Relative concentrations for the IP samples were determined from the standard curves for that sample and primer set. The on target relative concentration for each genotype was divided by either off target relative concentration to determine the enrichment of Gtf2ird1 binding.

### Hippocampus RNA-sequencing

#### Library preparation

The hippocampus was dissected from adult animals of the second behavior cohort (Table1). We used six animals of each genotype, three males and females of the WT and CD animals and two males and four females of the *Gtf2i\** genotype. The hippocampus was homogenized in 500ul of 1x RIPA supplemented with two RNAse inhibitors, RNAsin and SUPERase In, and 250ul of the homogenate was added to 750ul of Trizol LS and stored at −80° C until RNA extraction. RNA was extracted using the Zymo clean and concentrator-5 kit following the on column DNAse I digestion protocol and eluted in 30ul of water. The quality and concentration of the RNA was determined using a nanodrop 2000 and Agilent RNA Highsenstivity Tape screen ran on the TapeStation 2000 (Agilent). All RINe scores were above seven.

1ug of RNA was used as input and rRNA was depleted using the NEBNext rRNA Depletion kit (Human/Mouse/Rat). RNAseq libraries were prepared using the NEB Next Ultra II RNA library Prep Kit for Illumina. The final uniquely indexed libraries for each sample were amplified using the following thermocycler conditions: 98° C for 30 seconds, 98° C 10 seconds, 65° C 1 minute and 15 seconds, 65° C 5 minutes, hold at 4° C, repeat steps 2 through 3 6 times. Each sample had a unique index. Samples were pooled at equal molar amounts and 1×50 reads were sequenced on one lane of a HiSeq3000 at the Genome Technology Access Center at Washington University School of Medicine. The RNAseq data is available at GEO with accession number (submitted, waiting on accession number).

#### RNAseq analysis

The raw reads were trimmed of Illumina adapters and bases with base quality less than 25 using the Trimmomatic Software (64). The trimmed reads were aligned to the mm10 mouse genome using the default parameters of STARv2.6.1b (65). Samtools v1.9 (66) was used to sort and index the aligned reads. Htseq-count v0.9.1 (67) was used to count the number of reads that aligned to features in the Ensembl GRCm38 version 93 gtf file.

The htseq output was analyzed for differential gene expression using EdgeR v3.24 (68). Lowly expressed genes were defined as genes that had a cpm less than two across all samples. Lowly expressed genes were then filtered out of the dataset. We used the exactTest function to make pairwise comparisons between the three groups: WT versus *Gtf2i\**, WT versus CD, and CD versus *Gtf2i\**. Genes were considered differentially expressed if they had an FDR< 0.1.

GO analysis was performed using the goseq R package (69). Nominally significant up and down regulated genes for each comparison were considered differentially expressed genes and the background gene set included all expressed genes after filtering out the lowly expressed genes. The top 10 most significant go terms for each ontology category were reported. To test how unlikely it is to see these go terms given the differentially expressed genes from the genotype comparisons, we shuffled the genotypes among the samples and repeated the differential expression analysis and go term analysis 1000 times and counted how many times the same go terms were identified in the top ten most significant go terms.

### Behavioral tasks

#### Animal statement

All animal testing was done in accordance with the Washington University in St. Louis animal care committee regulations. Mice were same sex and group housed with mixed genotypes in standard mouse cages measuring 28.5 x 17.5 x 12cm with corn cob bedding and ad libitum access to food and water in a 12 hour light dark cycle, 6:00am-6:00pm light. The temperature of the colony rooms was maintained at 20-22° C and relative humidity at 50%. Two cohorts of mice were used in the behavior and RNA-seq experiments. The CD animals (Del (5Gtf2i-Fkbp6)1Vcam) were a gift from Dr. Victoria Campuzano and have been previously described (28) and were maintained on the C57BL/6J strain (https://www.jax.org/strain/000664). The first behavior cohort (Table 1) used *Gtf2i\** and CD females as breeders. The second behavior cohort (Table 1) used just CD female breeders as male CD animals were frequently not successful at breeding. Male and female mice were included in the behavior tasks. Experimenters were blind to genotype during all testing. Besides the maternal separation induced pup ultrasonic vocalization, all behaviors were done in adult animals older than 60 days and less than 150 days old. Mice were moved to the testing facility at least 30 minutes before the test to allow the mice to habituate to the room. The male experimenter was present during this habituation so the mice could also acclimate to the experimenter. Sex differences were assessed in all experiments, and are discussed when they were significant. Otherwise, the data is presented with the males and females pooled. Animals were removed from analysis if they were outliers, defined as having values greater than 3.5 standard deviations above or below the mean for their genotype group. Animals were also removed if the video and tracking quality were too poor to be analyzed. All filtering was conducted blind to genotype.

#### Maternal separation induced pup ultrasonic vocalization

To assess early communicative behaviors we performed maternal separation induced pup ultrasonic vocalization (USVs). Animals were recorded on postnatal day three and postnatal day five, days when FVB/AntJ animals begin to make the most calls (39). The parents were placed in a new cage, and the home cage containing the pups was placed in a warming box (Harvard Apparatus) set at 33° C for at least 10 minutes prior to the start of recording. Pups were individually placed in an empty standard-mouse cage (28.5 x 17.5 x 12cm) located in a MDF sound-attenuating box (Med Associates) that measures 36 x 64 x 60cm. Prior to recording, the pup’s skin temperature was recorded using a noncontact HDE Infrared Thermometer, as it has been shown that decreased body temperature elicits increased USVs (70). There was no difference in body temperature between genotypes (F_2,61_= 2.521, p=0.089)(Supplemental Table 1). USVs were detected using an Avisoft UltraSoundGate CM16 microphone placed 5cm above the bottom of the cage, Avisoft UltraSoundGate 416H amplifier, and Avisoft Recorder software (gain=3dB, 16bits, sampling rate =250kHz). Animals were recorded for 3 minutes, weighed, checked for detachment of pinnae, and then placed back into the home cage in the warming chamber. After all animals had been recorded the parents were returned to the home cage. Sonograms of the recordings were prepared in MATLAB (frequency range =25-120kHz, FFT [Fast Fourier Transform] size=512, overlap=50%, time resolution =1.024ms, frequency resolution = 488.2Hz) along with number of syllables and spectral features using a previously published protocol (39, 71) based on validated methods (72).

#### Sensorimotor battery

We assessed motoric initiation, balance, coordination, and strength as described in (73, 74) over two days using the following tasks: day 1) walking initiation, ledge, platform, pole; day 2) 60 screen, 90 screen, and inverted screen. Each task was performed once then the animals were allowed a 20 minute break then the tests were repeated in reverse order to control for practice effects. The two trials for each task were then averaged to be used in analysis. Walking initiation was tested by recording the time it takes for the mouse to exit a demarcated 24 x 24cm square on top of a flat surface. To assess balance, the mice were placed on a plexiglass ledge with a width of 0.5cm and a height of 38cm. We recorded how long the mouse balanced on the ledge up to 60 seconds. Another test of balance required the mouse to balance on a wooden platform measuring 3.0cm in diameter, 3.5cm thick and was 25.5cm high. The amount of time the animal balanced on the platform was recorded up to 60 seconds. Motor coordination was tested by placing the mouse at the top of a vertical pole with the head facing upward. The time it took the mouse to turn so the head was facing down was recorded as well as the time it took the mouse to reach the bottom of the pole up to 120 seconds. On day two the mice performed screen tasks that assessed coordination and strength. Mice were placed head facing downward in the center of a mesh wire grid that had 16 squares per 10cm and was 47cm off the ground and inclined at 60 degrees. The time it took the mice to turn and reach the top of the screen was recorded up to 60 seconds. Similarly the mice were placed in the center facing downward of mesh wire screen with 16 squares per 10cm, elevated 47cm from the surface of a utility cart, and inclined at 90 degrees. The time it took the mice to turn around and reach the top was recorded up to 60 seconds. To test strength, the mice were placed in the center of a mesh wire grid used for the 90 screen task and then inverted so the mouse was hanging from the screen that was elevated 47cm. The time the mouse was able to hang onto the screen up to 60 seconds was recorded.

#### One hour locomotor activity

We tested the animals’ general exploratory activity and emotionality in an one hour locomtor activity task (74). Animals were placed in the center of a standard rat cage (47.6 x 25.4 x 20.6cm) and allowed to explore the cage for one hour in a sound-attenuating enclosure with the lightening set to 24 lux. The one hour sessions were video recorded and the animals position and horizontal movements were tracked using the ANY-maze software (Stoelting Co.: RRID: SCR_014289). The apparatus was split into two zones: a 33 x 11cm center zone, and a bordering 5.5cm edge zone. ANY-maze recorded total distance traveled in the apparatus, and total distance traveled, time spent, and entries into each zone. The mouse was considered to have entered a zone when 80% of the body was detected within the zone. The rat cages are thoroughly cleaned with 70% ethanol between mice.

#### Marble burying

Marble burying is a task that measures the natural digging behavior of mice and is correlated to compulsive behaviors and hippocampal function (45). Following our previously published methods (74), a standard rate cage (47.6 x 25.4 x 20.6cm) was filled with autoclaved aspen bedding to a depth of 3cm and placed in a sound-attenuating enclosure with lighting set to 24 lux. 20 glass marbles were arranged in 5 x 4 grid on the surface of the bedding. Mice were placed in the center of the rat cage and allowed 30 minutes to explore and bury the marbles. The session was recorded using a digital camera and the animals horizontal movements and position in the apparatus were tracked using ANY-maze with the same center and edge zones as described in the one hour activity task. After 30 minutes mice were put back in their home cage and the number of marbles not buried was counted by two observers. A marble was considered buried if 2/3 of the marble was underneath the bedding. The average of the two scorers was used to calculate the average number of marbles buried. The marbles and rat cages were thoroughly cleaned with 70% ethanol between mice.

#### Three-chamber social approach

To assess voluntary sociability and preference for social novelty we used the three chamber social approach assay as previously described (74–76). The task took place in a plexiglass arena with two partitions with rectangular openings (5 x 8cm) dividing the arena into three chambers that each measure 19.5 x 39 x 22cm. The openings could be closed using plexiglass doors that slide into the openings. The task consisted of four consecutive 10 minute trials. During trial one the animals were habituated to the middle chamber with the openings to the side chambers closed. In trial two the animals were allowed to explore the entire apparatus. Trial three was the sociability trial. In one side chamber there was an empty steel pencil cup (Galaxy Pencil/Utility Cup, Spectrum Diversified Designs, Inc.) that was placed upside with an upside clear drinking cup secured to the top to prevent animals from climbing on top of the cup; this was the empty side. In the other side chamber there was an identical pencil cup that housed an age- and sex-matched, sexually naive, unfamiliar C57BL/6J stimulus animal; this was the social side. The pencil cups allowed sniffing behavior to occur and exchange of odor cues, but limited physical contact to prevent aggressive behaviors. The experimental animal was allowed to explore the whole apparatus. The side of the empty cup and social cup were counterbalanced across all the samples. In trial four we tested preference for social novelty. A new stranger stimulus animal was placed in the formerly empty cup. All stimulus animals were habituated to the apparatus and the cups for 10 minutes one day prior to testing. Each stimulus animal was used only once per day. During all trials the task was video recorded and the animal’s position, animal’s head, and movement was tracked with ANY-maze software. We quantified how much time the animal spent in each chamber, as well as distance traveled and number of entries. A 2cm area around the cups was defined as the investigation zone, and the animal’s head was used to determine when it was investigating the stimulus animals or the empty cup. The first five minutes of the social and novelty trials were analyzed because this is when the majority of the social investigation occurs (77). The entire apparatus was thoroughly cleaned after each animal using 2% chlorhexidine (Zoetis). The stimulus cups were cleaned using 70% ethanol.

#### Modified social approach

To test for habituation to social stimuli over extended amounts of time, we slightly modified the social approach task. We used the same apparatus as described above. We included an additional day of habituation to the apparatus for the experimental animals on the day prior to the actual test to ameliorate novelty induced exploration of the apparatus and to potentiate exploration of the investigation zones. During the habituation day the animals were placed in the center chamber for 10 minutes with the doors to the side chambers closed. Next, the animals were allowed to explore the whole apparatus for 20 minutes. The stimulus animals were habituated to the cups in the apparatus for 30 minutes prior to the test day. Trial one and trial two were the same as the social approach described above. For trial three, the sociability trial, the experimental animals were placed in a cylinder in the center chamber, while the empty cup and stimulus animal cup were being placed in the side chambers. This ensures a random starting direction for the experimental mouse so we could make an unbiased measure of which chamber the experimental mouse chose to enter first. The sociability trial lasted for 30 minutes, in which the experimental animal was allowed to freely explore the apparatus and investigate the empty cup and social cup. The social novelty trial was not conducted.

#### Tube test of social dominance

The tube test of social dominance tests for social hierarchy behaviors in mice (74, 78). This task took place over five days. Days one and two were habituation trials. During day one, the animals were placed in the left entrance of a clear acrylic tube measuring 3.6cm in diameter and 30cm in length and allowed to walk through the entire tube and exit the tube on the right side. Day two was the same but the mice started on the opposite side of the tube. These two habituation days allow the mice to acclimate to the tube, and potentiates task performance. On each of three consecutive test days, two mice of different genotypes were placed in the entrances to the tube and allowed to meet in the middle, at a clear acrylic partition. When both mice were at the acrylic partition, it was removed and the trial began. The trial ended when one mouse was pushed out or backed out of the tube so that all four paws were out of the tube, or two minutes had passed. The mouse that remained in the tube was considered the dominant winner and the mouse that was no longer in the tube was considered the submissive loser. If both mice were still in the tube after two minutes it was considered a tie. Each mouse was tested only once each day, and the mice were tested against a novel mouse each day. After each test, the tube was cleaned with 2% chlorhexidine (Zoetis) solution. All of the test sessions were recorded using a USB camera connected to a PC laptop (Lenovo). The observer scored the test from the videos.

#### Rotarod

The accelerating rotarod (Rotamex-5; Columbus Instruments, Columbus, OH) tests motor coordination, motor learning, and balance. We used a previously published rotarod paradigm (79–81) that tests animals on three conditions: 1) stationary rod 2) continuous rotation and 3) accelerating rotation during three different sessions that were separated by three days to minimize motor learning. During each day the animals had five trials; one stationary trial, two continuous trials, and two accelerating trials. During the stationary trial, the animals were placed on the stationary rod and the time that the animals stayed on the rod was recorded up to 60 seconds. During the continuous trials, the animals were placed on the rod rotating at three rotations per minute. The time the animals stayed on the rotating rod was recorded up to 60 seconds. In the accelerating trials, the animals were placed on the rod that was rotating at two rotations per minute. Once the animals were on the rotating rod, the rod began to accelerate at 0.1rpm and reached 17rpm at 180 seconds. The time the animals stayed on the rod up to 180 seconds was recorded. The two trials for the continuous rotation and accelerating rotation during each session were averaged for analysis. If an animal fell off the rod during any session within the first five seconds, the animal was placed back on the rod and the time was reset up to two times. If the mouse fell off within five seconds on the third try that time was recorded.

#### Elevated Plus Maze

The elevated plus maze was used to assess anxiety-like behaviors in mice using previously published protocols (76, 82, 83). The apparatus had two closed arms that measured 36 x 6.1 x 15cm, two open arms, and a central platform that measured 5.5 x 5.5cm. The time spent in the open arms was used as a measure of anxiety-like behavior in mice, since mice prefer to be in an enclosed area. Each mouse was tested once per day for three consecutive days. During the test the animals had five minutes to freely explore the apparatus. The animals position, movement, entries into each arm, and time spent in each arm were determined by beam breaks of pairs of photocells arranged in a 16 (x-axis) x 16 (y-axis) grid. Beam breaks were monitored by the Motor Monitor software (Kinder scientific). The test was conducted in the dark with black lights, and was recorded by an overhead digital camera using the night vision setting.

#### Pre-pulse inhibition (PPI)

To test for normal sensorimotor gating and normal acoustic startle response we performed PPI on the animals. Mice were placed in a cage located on top of a force transducer inside of a sound-attenuating box with a house light on (Kinder Scientific). The force transducer measured the startle response of the animals in Newtons. We used a protocol adapted from (76, 84). The protocol was run using the Startle Monitor II software (Kinder scientific). The protocol started with five minutes of acclimation to the 65dB background white noise, which is played continuously throughout the procedure. After acclimation there were 65 trials that pseudo-randomly alternated between different stimulus conditions, beginning with five consecutive trials of the startle stimulus, which was a 40msec 120dB pulse of white noise. The middle trials cycled through blocks of pre-pulse conditions, blocks of non-startle conditions, where only the background noise is played, and two blocks of startle conditions. Each block consisted of five trials. The testing ended with single trials of pulses played at 80dB, 90dB, 100dB, 110dB, followed by five more startle trials of 120dB. There were three different pre-pulse conditions, where a pulse of 4dB, 8dB, or 16dB white noise above the background sound was played 100msec preceding the 120dB startle stimulus. The average startle response during the middle two blocks of startle trials was considered to be the animal’s acoustic startle response(ASR). Each trial measured the startle of the animal for 65msec after the stimulus, and the average force in Newtons across this time was used as the startle response. The pre-pulse inhibition was calculated as the difference of the average ASR and the startle response during the respective pre-pulse trial (PP) divided by the ASR of the startle trials multiplied by 100: ((ASR – PP)/ASR)*100.

#### Contextual and Cued Fear Conditioning

Contextual and cued fear conditioning were used to assess associative learning and memory. We followed a previously published method (81, 85). The test occurred over three days. A camera placed above the apparatus recorded the session. Freezing behavior during each minute was detected in .75s intervals using the FreezeFrame (Actimetrics, Evanston, IL) software. Freezing behavior was defined as no movement except for normal respiration, and is presented as percent time freezing per minute. During day one, animals were allowed to explore the Plexiglas chamber (26cm x 18cm x 18cm; Med Associates Inc.) with a metal grid floor and a peppermint scent that was inaccessible to the animals. A trial light in the chamber turned on for the duration of the five minute trial. During the first two minutes animals were habituated to the apparatus, and freezing during this time was considered the baseline. An 80db white noise tone was played for 20 seconds at 100 seconds, 160 seconds, and 220 seconds during the test. During the last two seconds of the tone (conditioned stimulus CS) a 1.0mA foot shock (unconditioned stimulus UCS) was delivered. The mice were returned to their home cage at the end of the five minute trial. On day two contextual fear memory was tested. The animals were placed into the same chamber with peppermint scent and the illuminated light and no tone or shock was delivered. Freezing behavior was measured over the eight minute task. The amount of time freezing in the first two minutes on day two was compared to the baseline freezing on day one to test the effects of the contextual cues associated with the UCS from day one. On day threed the animals were placed in a new context, a chamber with black walls, and a partition that creates a triangle shaped area and an inaccessible coconut odor. During this 10 minute task, the trial light was on for the entire duration. The animals explored the apparatus for the first two minutes to determine baseline freezing and then the same 80dB (CS) tone from day one was played for eight minutes. The freezing behavior during this time tested the effects of the CS associated with the UCS shock from day one. Shock sensitivity was tested for each mouse three days after the cued fear test following the procedure previously described in (81). Mice were placed in the chamber with the wire grid floor and delivered a two second shock of 0.05mA. The mA of the shock was increased by 0.05mA up to 1.0mA. At each shock level the animal’s behavior was observed and the current level at which the animal flinched, exhibited escape behavior, and vocalized was recorded. Once the animal had exhibited each of the behaviors the test ended. Shock sensitivity assessment served to confirm differences in conditioned fear freezing were not confounded by differences in reactivity to the shock current.

#### Resident intruder

The resident-intruder paradigm, as described previously (86), was used as a direct social interaction test. Only males were used in this experiment. Male mice were individually housed in standard mouse cages for 10 days. Cages were not changed so the mice could establish a territory. The testing took place over three days in which the home cage of the experimental animal was placed in a sound-attenuating box in the dark with two infrared illuminators placed in the box. A clear Plexiglas covering with holes was placed over the cage to prevent animals from jumping out of the cage. A digital camera using the night vision setting recorded the task. On each day a WT C57BL/6J stimulus animal (intruder), age and sex matched was introduced into the experimental animal’s (resident) home cage. The animals were allowed to interact for 10 minutes after which the stimulus animal was removed from the cage. A stimulus animal was only used once per day. The testing was repeated for two more days, during which the experimental animals were paired with novel intruders.

The videos were tracked using Ethovision XT 13 software (Noldus Information Technology) using the social interaction module. This module allows for simultaneous tracking of two unmarked animals. The initial tracking was further corrected manually using the track editing tools, to ensure the head and the tail points were oriented correctly. All of the video tracks were smoothed first with the loess method and then with the minimal distance moved method. The variables of interest were the mean bout of time, frequency, and the cumulative duration of time that the experimental animal’s nose was less than 0.6cm from the stimulus animal’s nose, interpreted as nose-to-nose sniffing, or when the experimental animal’s nose was less than 0.45cm from the tail base of the stimulus animal, interpreted as anogenital sniffing. These distance thresholds were determined by an experimenter blind to genotype, examining the videos using the plot integrated view functionality to ensure that the events called by the software accurately defined the social behavior.

### Statistical Analysis

All statistical tests were performed in R v3.4.2. Western blots and qPCR were analyzed using a one factor ANOVA and the post hoc Tukey all pairwise comparison test was used determine differences between groups using the multcomp package (87).

For all behavior tests the data was assessed for univariate testing assumptions of normality and equal variances. Normality was assessed using the Shapiro-Wilkes test as well as manual inspection of qq plots. Equality of variances was tested using the Levene’s test. Behaviors that violated these assumptions were analyzed using non parametric tests. Repeated measures were analyzed using linear mixed models with the animal as the random effect. Significance of fixed effects were tested using the Anova function from the Car (88) package in R. Post hoc testing was done using the Tukey HSD test from the multcomp package. Tukey HSD test ‘within time point’ was used for post hoc repeated measures comparisons, as appropriate. See Supplemental Tables 1 and 6 for descriptions of all statistical tests.

## Acknowledgments

This work was supported by 1R01MH107515 (JDD), and the Autism Science Foundation, and the National Science Foundation Graduate Research Fellowship DGE-1745038 to NDK. We would also like to thank Dr. Victoria Campuzano for sharing the CD mouse line, and the Genome Technology Access Center for technical support, as well as Dr. Beth Kozel for critical advice on this project. We would also like to thank Dr. David Wozniak and the Animal Behavior Core at the Washington University School of Medicine for their time and resources. We would like to thank Rena Silverman for her contribution to the resident intruder analysis.

