## Supplemental Figure 1 for "*Gtf2i* and *Gtf2ird1* mutation are not sufficient to reproduce mouse phenotypes caused by the Williams Syndrome critical region"

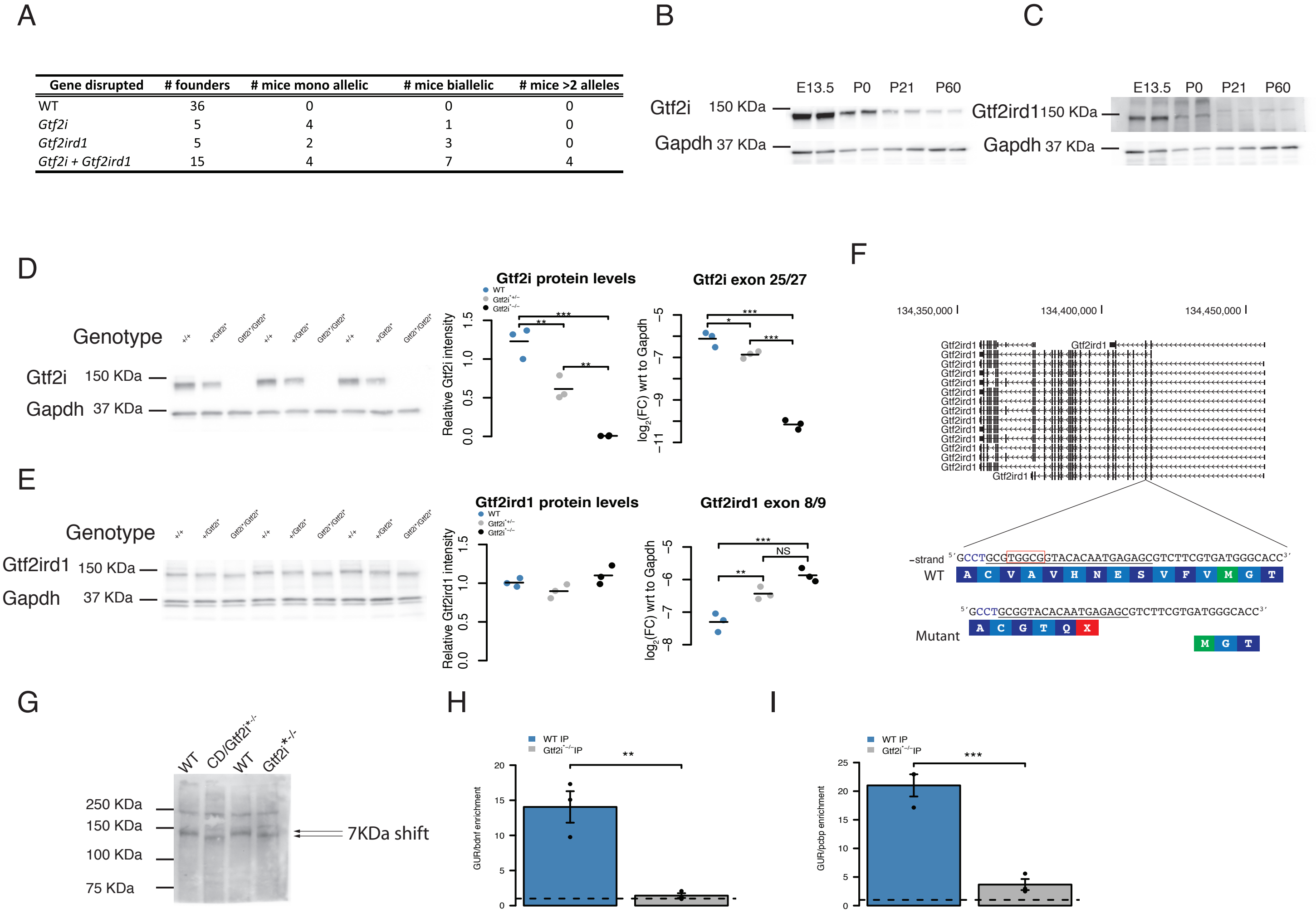

| Gene disrupted | # founders | # mice mono allelic | # mice biallelic | # mice >2 alleles |
|---|---|---|---|---|
| WT | 36 | 0 | 0 | 0 |
| *Gtf2i* | 5 | 4 | 1 | 0 |
| *Gtf2ird1* | 5 | 2 | 3 | 0 |
| *Gtf2i + Gtf2ird1* | 15 | 4 | 7 | 4 |



**Supplemental Figure 1. Generation of loss of function mutations in *Gtf2i* and *Gtf2ird1*. A** The number of founders from gRNA injection shows that two gRNAs are efficient at mutating both targets and have high rates of mosaicism. **B** Gtf2i protein is more highly expressed in the embryonic brain and is detectable in the adult brain, each time point includes two biological replicates. **C** Gtf2ird1 protein is more highly expessed in the embryonic brain and not detectable in the adult brain, each time point includes two biological replicates. **D** Gtf2i protein and transcript levels are decreased in the heterozyous *Gtf2i** mice and not detectable in the homozygous *Gtf2i** E13.5 brain. **E** Gtf2ird1 protein is not decreased in heterozygous or homozygous *Gtf2i** E13.5 brain, but the transcript is increased in heterozygous and homozygous animals. **F** Schematic of the consequences of the 5 bp deletion in *Gtf2ird1* showing the potential translation re-initation methionine in a new open reading frame. **G** A slight shift of Gtf2ird1 protein in animals homozygous and hemizgyous for the 5 bp deletion in exon 3 of *Gtf2ird1*, suggesting an N-terminal truncation of Gtf2ird1. **H** ChIP qPCR of the enrichment of the *Gtf2ird1* upstream regulatory sequence (GUR) over an off target sequence 7kbp upstream of *Bdnf* transcription start site in WT versus *Gtf2i** homozygous E13.5 brain. **I** ChIP qPCR of the enrichment of the *Gtf2ird1* upstream regulatory sequence (GUR) over an off target sequence 10kbp upstream of *Pcbp3* transcription start site in WT versus *Gtf2i** homozygous E13.5 brain. * $p < 0.05$, ** $p < 0.01$, *** $p < 0.001$
