## Supplemental Figure 2 for "*Gtf2i* and *Gtf2ird1* mutation are not sufficient to reproduce mouse phenotypes caused by the Williams Syndrome critical region"

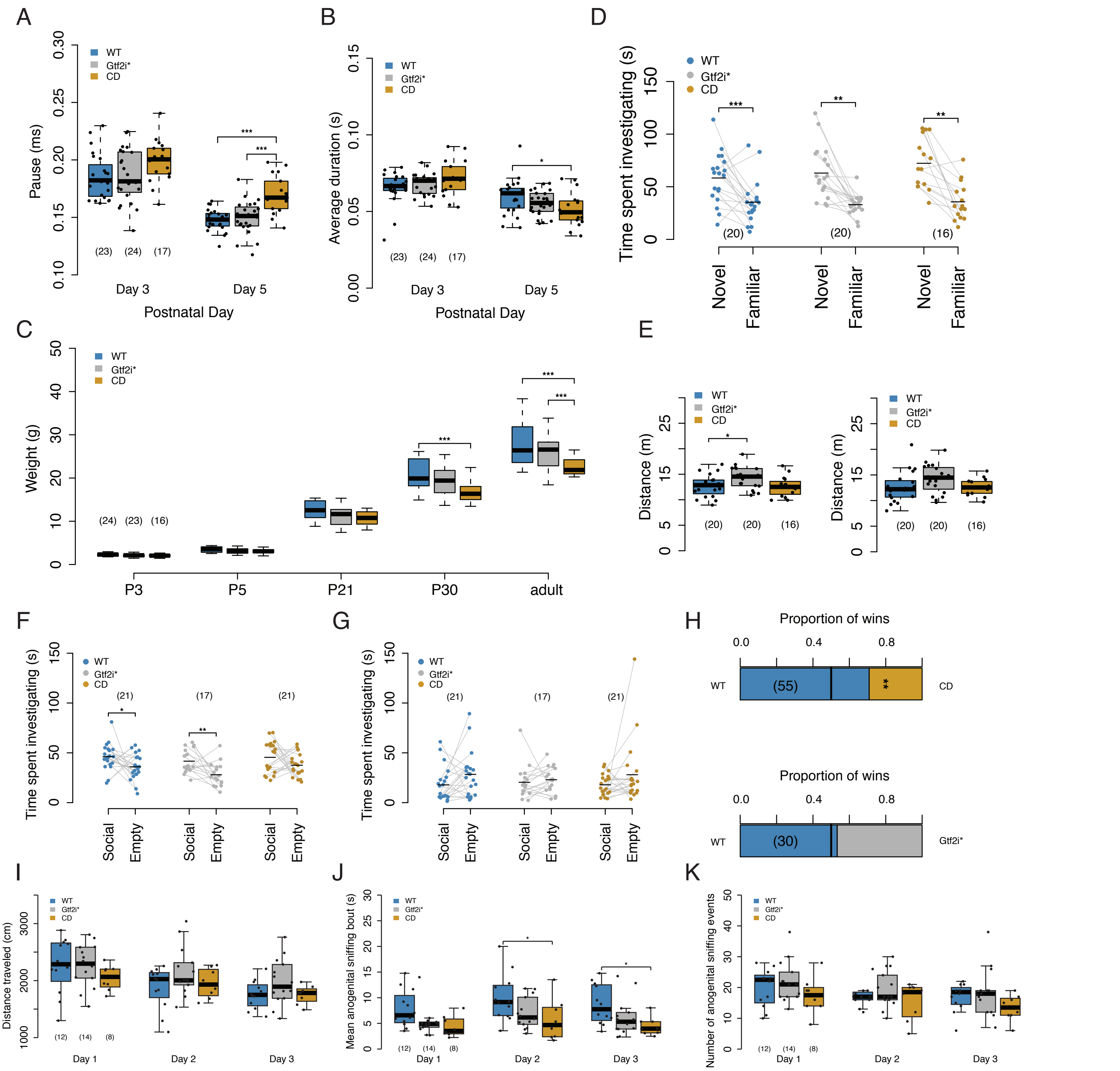


**Supplemental Figure 2. Social behaviors in CD and *Gtf2i** mutants. A** CD animals have increased pauses between bouts of USVs. **B** CD animals have decreased duration of USVs. **C** CD animals have decreased weight in adulthood, developmental weight does not explain differences in USV. **D** All genotypes show preference for social novelty. **E** Double mutants show increased activity in the social approach and social novelty trials of three chambered social appraoch. **F** WT and double mutants show social preference in the first 5 minutes of the extended social approach, but the CD mice are trending. **G** None of the genotypes show preference for social stimulus during the last 5 minutes of the extended social approach. **H** CD mice on C57BL6/J background show a submissive phenotype in tube test of social domiance while the double mutants show no phenotype on FVB/ANTJ background. **I** All genotypes travel similar distance in the resident intruder task. **J** CD animals have decreased mean bout time of anogenital sniffing in the resident intruder task. **K** However all genotypes have similar frequencies of anogential sniffing.
* $p < 0.05$, ** $p < 0.01$, *** $p < 0.0001$. Sample sizes are shown as numbers in parentheses
