## Supplemental Figure 3 for "*Gtf2i* and *Gtf2ird1* mutation are not sufficient to reproduce mouse phenotypes caused by the Williams Syndrome critical region"

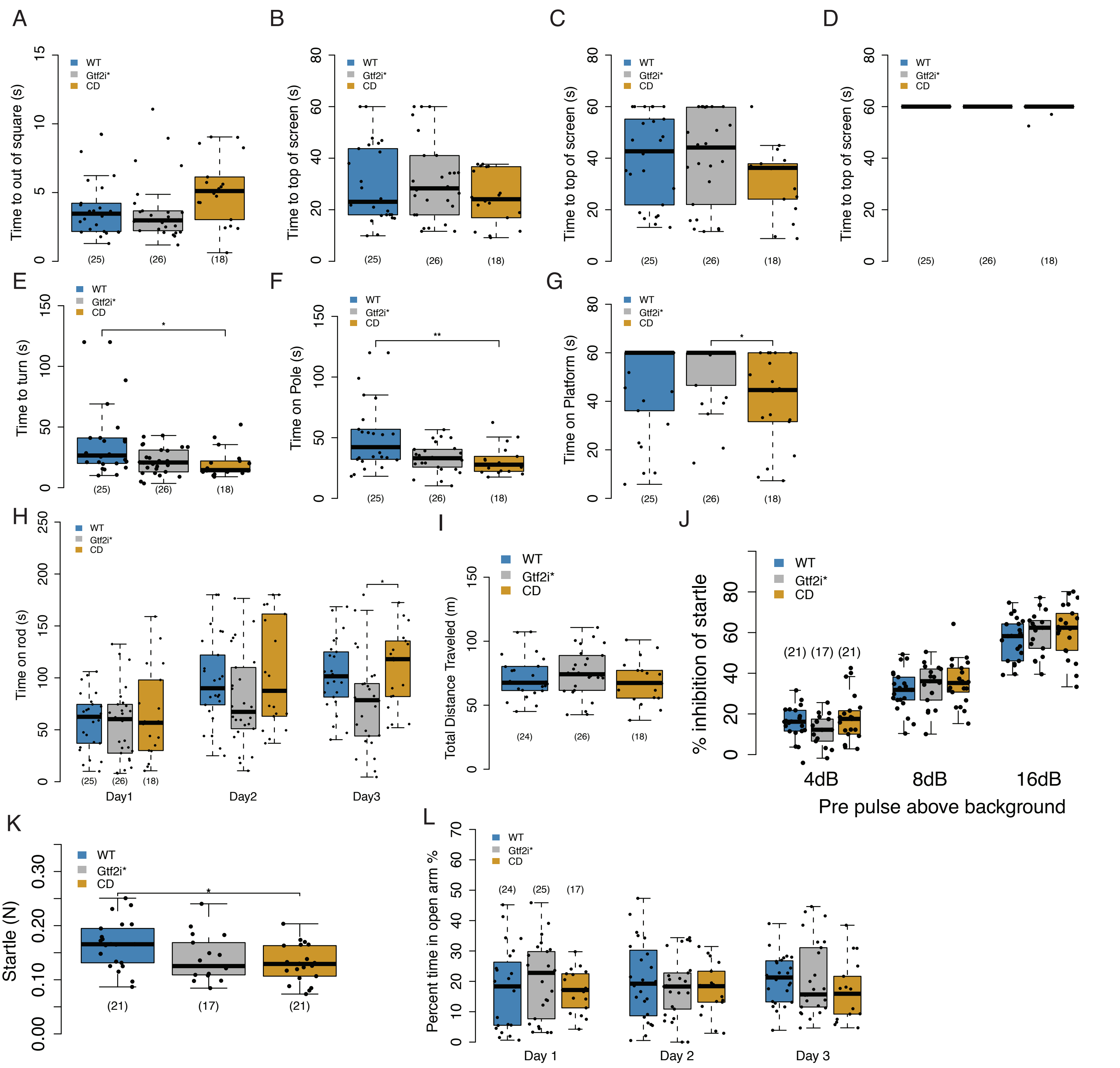


**Supplemental Figure 3. Motor and anxiety phenotypes in double mutants and CD animals. A** All animals show similar time to initiate walking. **B** All animals reach the top of a 60 degree inverted screen in similar amounts of time. **C** All animals reach the top of a 90 degree inverted screen in similar amounts of time. **D** All animals can hang onto an inverted screen for similar amounts of time. **E** CD animals are able to turn their bodies 180 degrees on a pole quicker than WT animals. **F** CD animals are able to reach the bottom of a pole quicker than WT littermates. **G** CD animals tend to fall off a platform more than double mutants. **H** On day 3 of the rotorod task double mutants fall off sooner than the CD animals. **I** All genotypes travel similar total distances in the marble burying assay **J** All genotypes show normal PPI. **K** CD animals have decreased startle to 120dB stimulus overall but this is due to decreased weight. **L** All genotypes spend similar amoutns of time in the open arm during elevated plus maze. * $p< 0.05$, ** $p < 0.01$, *** $p < 0.0001$. Sample sizes are shown as numbers in parentheses
