## Supplemental Figure 4 for "*Gtf2i* and *Gtf2ird1* mutation are not sufficient to reproduce mouse phenotypes caused by the Williams Syndrome critical region"

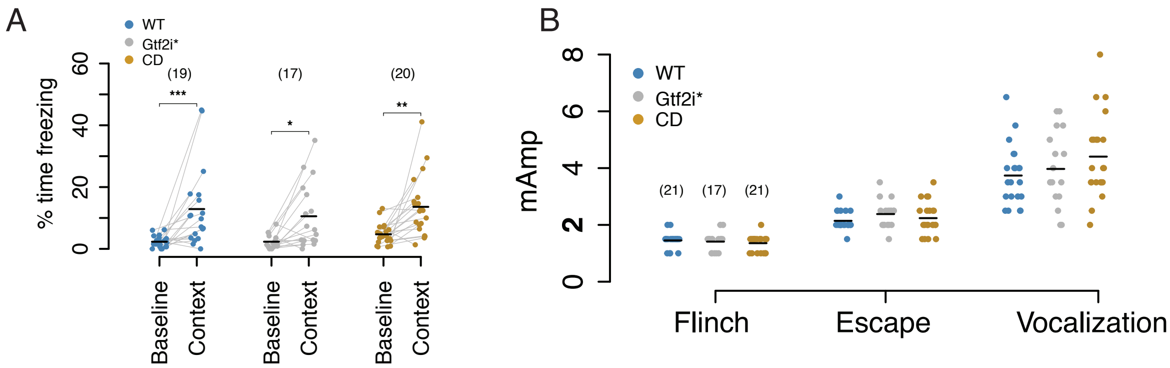


**Supplemental Figure 4. Contextual fear and shock senstivity in WS mutant models.** **A** All genotypes show a contextual fear response. **B** The response to foot shock is similar across all genotypes. * $p < 0.05$, ** $p < 0.01$, *** $p < 0.001$ Sample sizes are shown as numbers in parentheses
