## Supplemental Figure 5 for "*Gtf2i* and *Gtf2ird1* mutation are not sufficient to reproduce mouse phenotypes caused by the Williams Syndrome critical region"

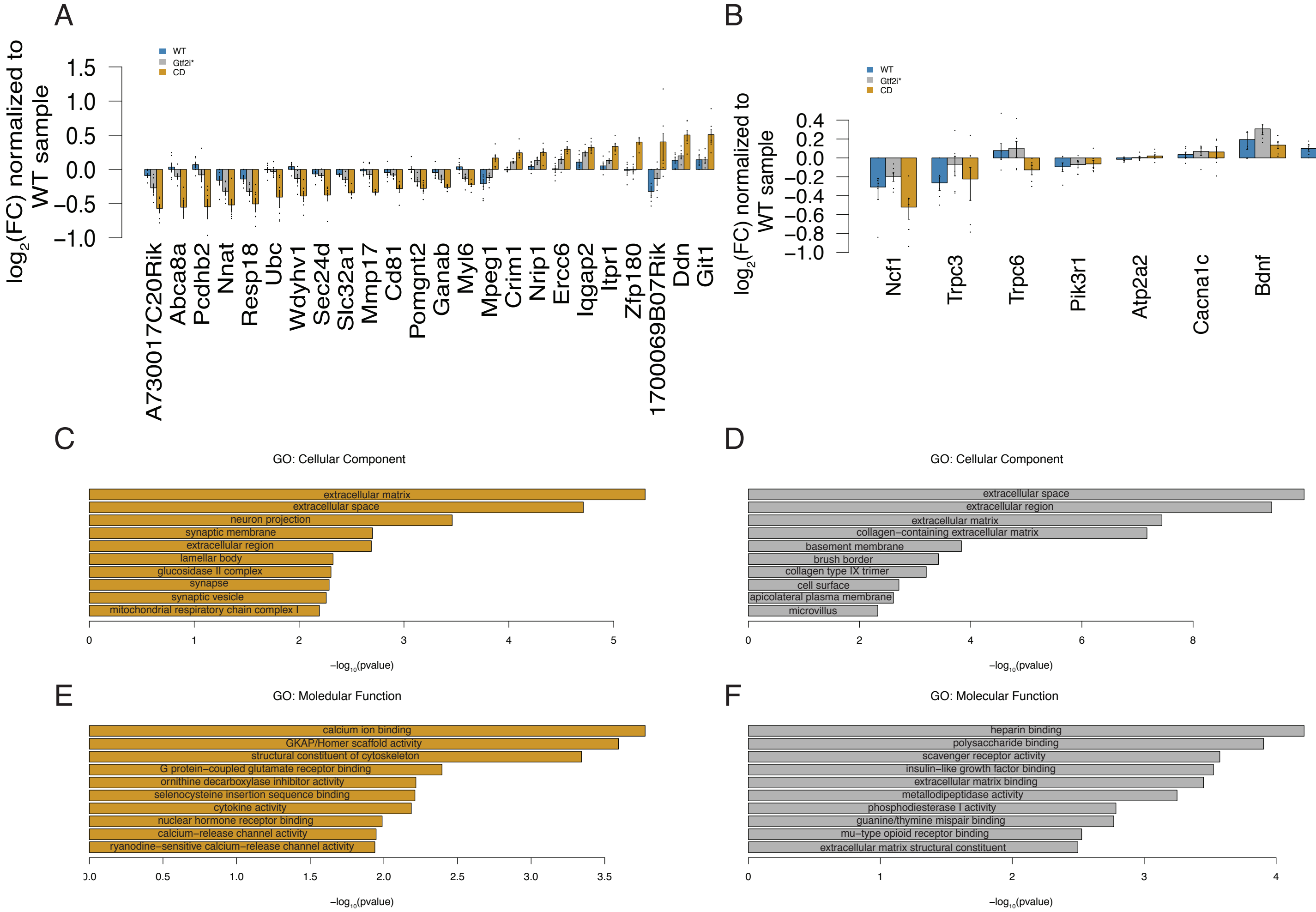


**Supplemental Figure 5. Small changes in hippocampal transcriptomes of WS models.** **A** Fold change of differentially expressed genes between WT and CD animals at an FDR < 0.1 normalized to WT levels. **B** Fold change of genes previously tested in CD hippocampus RNAseq from Ortiz-Romero et al. 2018. **C** The top ten enriched Cellular Component gene ontologies for genes that are nominally up or down regulated between CD and WT animals. **D** The top ten enriched Cellular Component gene ontologies for genes that are nominaly up or down regulated between *Gtf2i** and WT animals. **E** The top ten enriched Molecular Function gene ontologies for genes that are nominally up or down regulated between CD and WT animals. **F** The top ten enriched Molecular Function gene ontologies for genes that are nonminally up or down regulated between *Gtf2i** and WT animals
